# How a reaction-diffusion signal can control spinal cord regeneration in axolotls: A modelling study

**DOI:** 10.1101/2023.08.21.554065

**Authors:** Valeria Caliaro, Diane Peurichard, Osvaldo Chara

## Abstract

Axolotls are uniquely able to completely regenerate the spinal cord after amputation. The underlying governing mechanisms of this regenerative response have not yet been fully elucidated. We previously found that spinal cord regeneration is mainly driven by cell cycle acceleration of ependymal cells, recruited by a hypothetical signal propagating from the injury. However, the nature of the signal and its propagation remain unknown. In this theoretical study, we investigated whether the regeneration-inducing signal can follow a reaction-diffusion process. We developed a computational model, validated it with experimental data and showed that the signal dynamics can be understood in terms of reaction-diffusion mechanism. By developing a theory of the regenerating outgrowth in the limit of fast reaction-diffusion, we demonstrate that control of regenerative response solely relies on cell-to-signal sensitivity and the signal reaction-diffusion characteristic length. This study lays foundations for further identification of the signal controlling regeneration of the spinal cord.

## 1 Introduction

In contrast to most vertebrates, salamanders are capable of remarkable regeneration traits. Although more than 250 years have passed since the original discovery of salamander tail regeneration after amputation by Spallanzani^1^, the governing mechanisms underlying these unparalleled regeneration capabilities have not yet been completely elucidated. The axolotl (*Ambystoma mexicanum*) is a paedomorphic salamander which can resolve severe and extreme injuries of the spinal cord throughout complete and faithful regeneration^2,3,4^.

Tail amputation in the axolotl triggers the reactivation of a developmental-like program in the ependymal cells, neural stem cells lining in the central canal of the spinal cord^5^. Although cell influx, cell rearrangements, and re-entrance to the cell cycle from quiescence can be observed, the main driver of the spinal cord outgrowth is the acceleration of the cell cycle^6^. Indeed, during regeneration, ependymal cells anterior to the amputation plane shorten their cycle length three times^5^. We previously identified a high-proliferation zone emerging 4 days after amputation within the 800 *μm* anterior to the amputation plane that shifts posteriorly during spinal cord regeneration^6^. This particular spatiotemporal pattern was quantified with a recruitment limit curve separating the high-proliferation from the low-proliferation zones^6^. We recently developed the first 1D cell-based computational model of the axolotl spinal cord in which each ependymal cell was simulated with an internal clock portraying its age/position along its cell cycle^7^. The model assumed that tail amputation triggers a hypothetical signal that propagates anteriorly with constant velocity for a certain time along the spinal cord, supposedly one-dimensional, recruiting ependymal cells when the signal concentration is higher than zero. By adapting the Fluorescent Ubiquitination-based Cell Cycle Indicator (FUCCI) technology to axolotls (AxFUCCI), we were able to visualize cell cycles *in vivo*, qualitatively reproducing the model predicted distribution of ependymal cells in coordinates of time, space and cell cycle^7^. Nevertheless, the signal and its nature remain to be elucidated.

Numerous signalling pathways control morphogenetic processes whose dynamics operate under a reaction-diffusion mechanism during development^8,9,10,11^ and regeneration^12,13^. A very well-known example consists of one morphogenetic signal propagating through the tissue while being subjected to an enzymatic degradation or cellular uptake / elimination^14^. This last process constitutes the “reaction” responsible for the consumption of the signal and the consequent reduction in signal concentration. In this study, we aim to determine whether this mechanism could explain the regenerative response observed in the axolotl spinal cord by following a modelling approach combining computational modelling with theory.

To that aim, we here propose a hybrid multi-scale cell-based computational model of the axolotl spinal cord combining the ependymal cell layer with a signal that follows a reaction-diffusion scheme while orchestrating the regenerative response by accelerating the ependymal cell cycle. The model successfully fits the aforementioned recruitment limit curve^6^ and correctly predicts the spinal cord outgrowth^5^, providing a first estimate of the diffusion coefficient and half-life of the hypothetical regeneration-inducing signal. By using a subsequent rigorous theoretical approach, we prove that the spinal cord growth emerging during regeneration in the axolotl can be controlled by the reaction-diffusion characteristic length and the ependymal cell-to-signal sensitivity in the regime of fast diffusion and reaction. Finally, we further corroborated the estimations of signal diffusion coefficient and half-life by comparing the computational model simulations with the experimental spatiotemporal distribution of cells in G1/G0 and S/G2 cells extracted from AxFUCCI axolotl spinal cords during regeneration. Ultimately, this study provides insights into the biophysical properties of signalling processes responsible for successful axolotl spinal cord regeneration.

## 2 Results

### 2.1 Hybrid multi-scale model of the ependymal tube controlled by a reaction-diffusion signal qualitatively reproduces spinal cord regeneration in the axolotl

To assess whether spinal cord regeneration in the axolotl can be explained by a potential signaling mechanism operating under a reaction-diffusion scheme, we developed a computational model of the regenerating spinal cord where the ependymal cell division rate is under the control of a generic signal. Specifically, we proposed a multi-scale and hybrid model: while the ependymal cells are featured as non-overlapping and proliferating rigid discs in a “cellular” scale, the density of the signal controlling the ependymal cells is represented by a continuous field in the “signaling” scale (Fig. 1 A, B).

**Figure 1:**
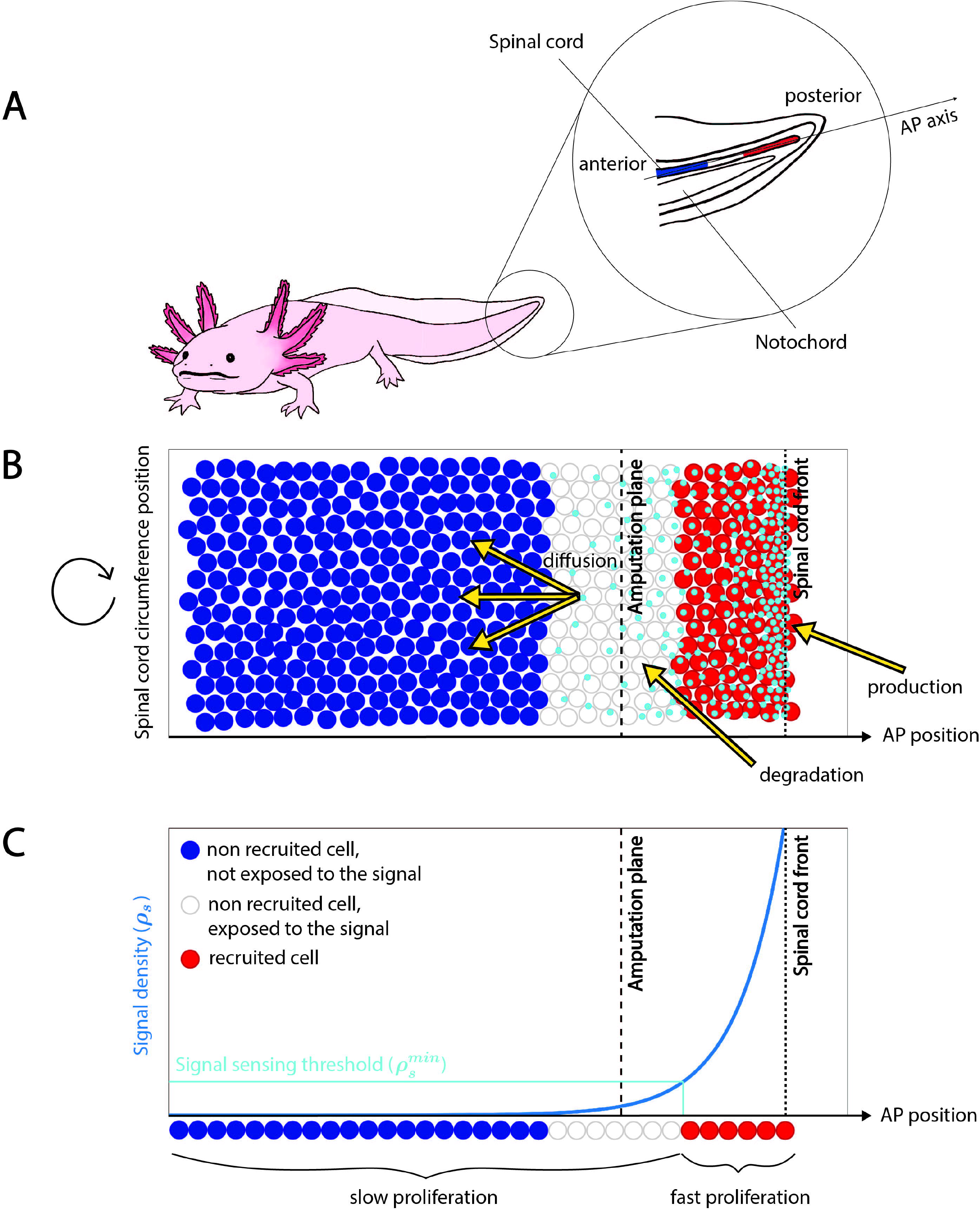
Schematic representation of the computational model of the signal-dependent regeneration of axolotl spinal cord. **(A)** Sketch of the axolotl showing the regenerating spinal cord in the tail. **(B)** Scheme of the model components. The y-axis represents position along the spinal cord circumference and the x-axis corresponds to the Anterior-Posterior axis. The ependymal cells are represented as discs coloured with a French flag-like code, depending on whether they are not exposed to the signal and are not recruited (blue), or they are exposed to the signal but are not recruited (white), or they are exposed to the signal and are recruited (red). The signal is represented as cyan particles that are produced at the front of the tissue, have a finite half-life (degradation) and diffuse in the domain populated by ependymal cells. **(C)** Cell recruitment depends on the local signal density (whose density is here represented in a 1D view). A cell is recruited when it is exposed to a signal density that overcomes the threshold 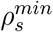 (indicated in cyan). The x-axis represents the AP axis, the blue curve is the density of the signal. In (B) and (C), the amputation plane is indicated by a dashed black line and the spinal cord front is indicated by a dotted black line.

We assumed a 2D domain for both scales, where the two spatial dimensions correspond to the Anterior-Posterior (AP) axis and the perpendicular direction given by the arc length along the circumference of the central canal apical surface (Circumference axis, Fig. 1. A). We modeled the reaction-diffusion mechanism for the signal by assuming that the signal density *ρ*_*s*_(*x, t*) linearly degrades with a degradation rate *k* (which is inversely proportional to the half-life *τ*) and diffuses with diffusion coefficient *D* in the domain occupied by the ependymal cells. Mathematically, this can be summarized as follows:

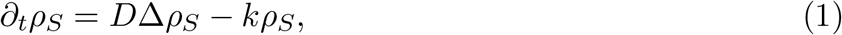

where the signal density is assumed equal to a non-zero constant at the posterior front of the spinal cord, at any given time *t*, modeling the secretory signalling centre of the wound epider-mis (see details of the boundary conditions for both scales in STAR methods section). Note that although reaction-diffusion models are typically composed of two or more species capable of Turing patterns^15,16,17,18^, Eq. (1) describes the simplest reaction-diffusion equation one can consider, in which the signal diffuses while undergoing homogeneous degradation within the tissue. We numerically solved the signaling dynamics by using the Smoothed-Particle Hydrodynamics (SPH) method^19^ which approximates the density function *ρ*_*S*_(*x, t*) by a cloud of *N* “signalling” particles of equal and constant masses, convected along their regularized concentration gradient following the diffusion velocity method^20^ (see justification of this method as well as details on the numerical implementation in STAR method section).

Based on our previous study^7^, we assumed that in the absence of the signal, the ependymal cells progress along their cell cycle with a certain pace (essentially, the cell cycle length is supposed lognormally distributed and the initial position of the cells within the cell cycle follows an exponential distribution; for more details of the stochastic component of the model, see^7^). Cells exposure to a high signal density leads to cell recruitment, manifested by G1 and S phase shortening and, ultimately, reduction of cell cycle length. In contrast to our previous article, we now assume that the recruitment sensitivity *S*_*R*_ is finite (defined as the inverse of the minimal signal density required to recruit an ependymal cell, 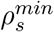, Fig. 1 C). We arbitrarily considered that cells are not exposed to the signal if there are less than two signalling particles around them. We supposed that cell recruitment is irreversible and inherited by the daughter cells. Thus, in our model, spinal cord growth during regeneration is driven by ependymal cell proliferation, in turn controlled by the signal diffusing from the spinal cord regenerating tip towards the anterior side while irreversibly recruiting new ependymal cells. We first validated the cellular scale of the model by by comparing our model results with a previous 1D model^21^ (Supplementary section 2.1, Fig. S1 C). Next, we validated the numerical scheme of the reaction-diffusion process governing the signaling scale of our model by comparing the 1D density profile of the signal with the analytical solution of the reaction-diffusion model in a finite domain (without ependymal cell proliferation), derived following our method previously reported^21^ (Supplementary section 2.2, Supplementary Fig. S1 D,E,F). After both scales of the model were validated, we evaluated whether the full model could generate a dynamical behavior consistent with the regenerative response observed in the axolotl spinal cord after amputation. Our simulations showed that the spatial distribution of the signal is shifted posteriorly as the spinal cord expands, result qualitatively equivalent to that previously reported^7^ (Fig. 2 A, B, Supplementary Movie 1).

**Figure 2:**
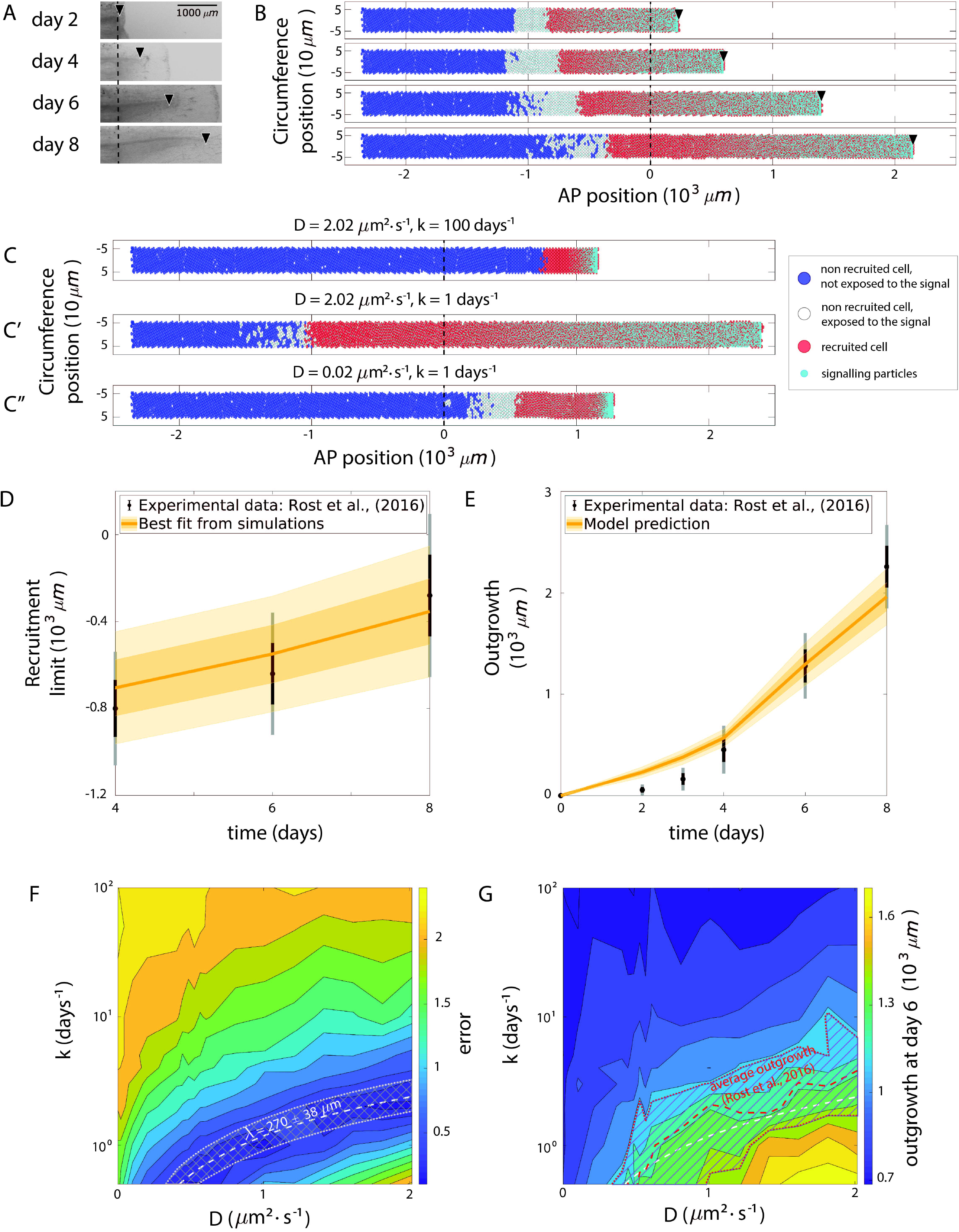
Axolotl spinal cord regeneration can be qualitatively and quantitatively explained by a signal operating under a reaction-diffusion regime. **(A)** Images of a regenerating spinal cord upon tail amputation at different times post amputation(modified from^6^, Fig. 1A). Scale bar, 1000 *μm*. **(B)** Simulations of the modelled regenerating spinal cord after tail amputation at the indicated times. In (A, B), the black dashed line represents the amputation plane while the black arrowheads indicate the tip of the regenerating spinal cord. Ependymal cells are represented as discs and colored using a French flag-like code, as explained in the legend of Fig. 1. B,C. Smaller cyan discs represent the signalling particles. Signal diffusion coefficient *D* = 1 *μm*^2^ *· s*^−1^, degradation rate *k* = 1 days^−1^ and 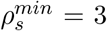. **(C,C’,C”)** Simulation results at day 8 post amputation: large diffusion and degradation of the signal (C), large diffusion and small degradation of the signal (C’), small diffusion and degradation of the signal (C”). Same representation as in panel (B). **(D)** Model-predicted time course of recruitment limit (yellow) fitted to the experimental counterpart^6^ (black). **(E)** Model-predicted time evolution of the spinal cord outgrowth (yellow) and the experimental counterpart^6^ (black). (D) and (E): for the model predictions, means are represented as solid lines and each shaded area corresponds to 1 standard deviation out of the best fitting simulations identified following the procedure described in STAR Methods); for the experimental data, means are represented by points while error bars represent the standard deviation of the recruitment limit posterior distribution in (D) and the standard deviation of the outgrowth measurements in (E), both obtained from the experiments reported in^6^. Black error bars correspond to 1 *σ* while grey error bars correspond to 2*σ*. **(F)** Phase portrait depicting the error between the model-predicted recruitment limit and the experimentally measured one^6^ at day 6, as a function of the diffusion coefficient *D* (x-axis) and the degradation rate *k* (y-axis), with the curve 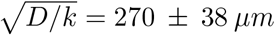 superimposed. **(G)** Phase portrait showing the tissue outgrowth predicted by the model at day 6 post amputation as a function of the diffusion coefficient *D* (x-axis) and the degradation rate *k* (y-axis), with the experimental outgrowth from^6^ (mean is represented by the red dashed line, the striped area correspo<u>nds</u> to 1 standard deviation, obtained from the experiments as reported in^6^) and the curve 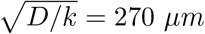 (white dashed line) superimposed.

### 2.2 Axolotl spinal cord regeneration can be quantitatively explained by a signal operating under a reaction-diffusion regime

By exploring the model parameters space we observed that the evolution of spinal cord outgrowth is controlled by the interplay between the diffusion coefficient *D* and the degradation constant *k* (Fig. 2 C, C’ and C”, Supplementary Movie 2). Indeed, decreasing the degradation constant (for a fixed diffusion coefficient value) is followed by an increase of the tissue outgrowth (compare Fig. 2 C and C’). On the contrary, decreasing the diffusion speed (while fixing the degradation) led to a remarkable decrease of the tissue outgrowth (compare Fig. 2 C’ and C”). Thus, there is an optimal balance between the diffusion of the signal (that governs the speed of the signal spreading and therefore the recruitment of new cells) and its degradation constant (that prevents the signal to travel too far during its half-life) that shapes the simulated regenerative response.

To gain a quantitative understanding of the regenerative response of the axolotl spinal cord, we fully parametrized the model. To that aim, we fixed the model parameters controlling the cell cycle paces and cellular geometry by using our previously reported experimental data obtained from ependymal cells in regenerating and uninjured spinal cords^5^ (see Table S1). Since the signal here proposed is hypothetical, we considered *D, k* and *S*_*R*_ as free parameters and explored the resulting model parameters space.

To assess if the model could explain the regenerative response of the axolotl spinal cord and to simultaneously estimate the free parameters, we fitted the model to the previously reported experimental recruitment limit, defined as the spatial position (in the AP axis) separating the (posterior) high-proliferation region from the (anterior) low-proliferation region^6^ (for details of the fitting procedure, see STAR methods). To this aim, we tracked the AP-axis position of the most anteriorly recruited cell in our simulations, denoted as *ξ*(*t*), and referred to it as the theoretical recruitment limit^7^. The model-predicted recruitment limit successfully reproduced the experimental switchpoint in a region along the curve 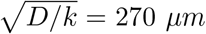 (Fig. 2 D). Importantly, this parametrization leading to the best-fitting of the switchpoint curve quantitatively predicted the time course of tissue outgrowth observed *in vivo*^6^ (Fig. 2 E). We found that there is a zone within the parameter space that allows the experimental switchpoint curve to be recovered (Fig. 2 F). This zone remarkably recapitulates the experimental outgrowth at day 6 (Fig. 2 G). Indeed, the distance between the experimental and the simulated recruitment limit curves (denoted by *E*(***ξ, C***_***e***_) and defined in Eq. (12) in STAR Methods) is minimal in a region along the curve 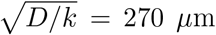 (Fig. 2 F). In this zone of the parameters space, we find the best agreement between the experimental spinal cord outgrowth at day 6 and the simulated outgrowth predicted by our model (Fig. 2 G). We emphasise that the model was not fitted directly to the experimental outgrowth quantified during axolotl spinal cord regeneration, but rather to the experimental recruitment limit dataset, and the best-fitting model prediction of tissue growth was then compared with experimental spinal cord outgrowth (Fig. 2 panels D and F). Altogether, these results suggest that spinal cord regeneration could be mainly controlled by a signal created at the (moving) boundary of the tissue while evolving via a reaction-diffusion process and recruiting cells as it spreads over the spinal cord. New cells are locally activated by the presence of this signal, accelerating tissue growth. In this scenario, the speed of tissue expansion is mainly controlled by the interplay between diffusion and degradation of the signal via the quantity 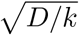. This quantity is well known in developmental biology in the cases of exponential morphogen gradient formation and referred to as the Characteristic Length *λ*^14^. It corresponds to the distance between the signal source (in our case, the posterior border of the tissue) and the position at which a concentration profile exponentially decaying equals a fraction 1*/e* of its concentration at the source.

### 2.3 The regenerative response of the spinal cord can be modulated by the cell sensitivity to the signal

In the previous section, we assumed a relatively small value of 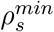 which results on extreme sensitivity of the ependymal cells to the signal density. Higher values of this parameter reduce the cell sensitivity to the signal (Supp. Fig. S2 A). In this section we investigated the role of the cell to signal sensitivity on the emergent regenerative outgrowth, by exploring the values of the parameter 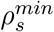 in the range indicated in Table S1. As expected, decreasing the sensitivity of cells to the signal leads to a reduction in the recruitment limit that reduces the spinal cord outgrowth (Fig. 3 A, A’, Supplementary Movie 3). The pool of recruited cells that activate their high proliferation program is directly linked to the ability of individual cells to measure the signal density around them. This result shows that the main component of tissue regeneration lies in the pool of recruited ependymal cells, the size of which is controlled by the signal properties and the cells sensitivity to the signal.

**Figure 3:**
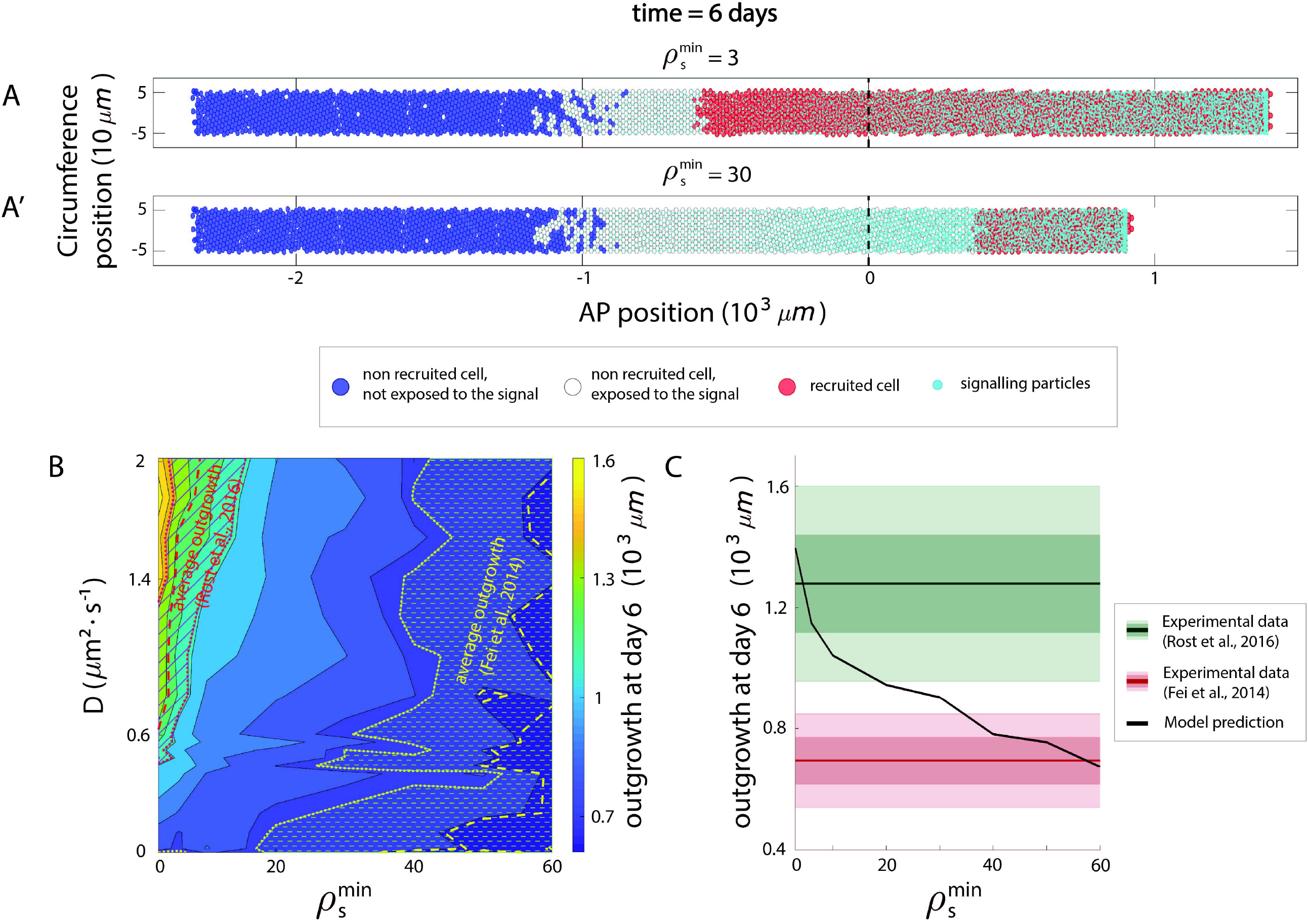
The regenerative response of the spinal cord can be modulated by the cell sensitivity to the signal. **(A, A’)** High cell-to-signal sensitivity (small 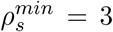, A) results in a higher recruitment limit (the domain of red recruited cells and outgrowth) compared to low cell-to-signal sensitivity (large 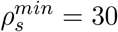, B). In (A, A’), model simulation results of the regenerating spinal cord at day 6 post amputation. Ependymal cells are represented as discs and colored using a French flag-like code, as explained in the legend of Fig. 1 B,C. **(B)** Phase portrait showing the model-predicted spinal cord outgrowth at day 6 for *k* = 1 days^−1^ as a function of the cell-to-signal sensitivity 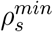 (x-axis) and the signal diffusion coefficient *D* (y-axis), with experimental measurements of^6^ (red) and^22^ (yellow) superimposed. **(C)** Spinal cord outgrowth at day 6 post amputation as a function of the cell-to-signal sensitivity 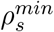. Decreasing the cell-to-signal sensitivity reduces the spinal cord outgrowth predicted at time 6 post amputation from values consistent with normal regenerative conditions^6^ (green) to those corresponding to impeded regeneration by knocking out SOX2^22^ (red). Means are represented as lines and each shadowed area corresponds to 1 standard deviation. In (A, C), signal diffusion coefficient *D* = 1 *μm*^2^ *· s*^−1^ and degradation rate *k* = 1 days^−1^.

We set out to determine quantitatively the influence of the ependymal cells sensitivity to the signal on spinal cord regeneration in the axolotl. To achieve this, we observed the outgrowth predicted by the model at day 6 post amputation, within the surface of the parameters space corresponding to the diffusion coefficient and the cell-to-signal sensitivity (Fig. 3 B and Supp. Fig. S2 B). We observed that decreasing the cell-to-signal sensitivity (by increasing 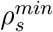) from the best-fitting value (that coincides with the experimental response previously observed at that time^6^) leads to the reduction of the outgrowth predicted at day 6 (Fig. 3 B and C). Interestingly, the model predicts that reducing the cell-to-signal sensitivity in 92.5% leads to a spinal cord outgrowth in agreement with that measured in the axolotl when the neural stem cell marker Sox2 is knocked-out^22^ (Fig. 3 B, C). Thus, our computational model of the axolotl spinal cord governed by cell cycle acceleration, in turn controlled by a reaction-diffusion signal, was validated by fitting to the previously reported experimental recruitment limit curve and correctly predicted spinal cord outgrowth observed under regenerative conditions. Importantly, our model gives a conceptual framework to interpret the lack of regenerative response of the Sox2 knock-out axolotls by turning down the sensitivity of the ependymal cells to the signal.

### 2.4 The control of spinal cord regeneration lies in the characteristic length of the signal and the cell-to-signal sensitivity

The previous sections showed that spinal cord regeneration in the axolotl can be quantitatively explained in terms of cell cycle acceleration of ependymal cells, controlled by a signal operating under a reaction-diffusion mechanism. This could imply that the control of the regenerative response relies on the three model parameters, the diffusion coefficient, the degradation constant/half-life of the signal and the cell-to-signal sensitivity. Nevertheless, visual inspection of Fig. 2 F, G depicts a clear domain identified by the characteristic length *λ* that recapitulates both the experimental switchpoint and the outgrowth at day 6. This suggests that the regenerative response is not controlled by the diffusion coefficient, half-life of the signal and cell-to-signal sensitivity, but rather by this sensitivity and *λ*. To test this hypothesis, we used our computational model to predict the regenerative response at a fixed time (We have arbitrarily chosen day 8) for different combinations of signal-related parameters and plotted the spinal cord outgrowth as a function of the characteristic length (Fig. 4 A). Interestingly, our results showed that the tissue outgrowth is almost an affine function of *λ* (*i*.*e*., showing a linear correlation with *λ*), the slope of which depends on the cell-to-signal sensitivity. This result indicates that the control of the regenerative response falls on the signal characteristic length and the sensitivity of the cells to the signal.

**Figure 4:**
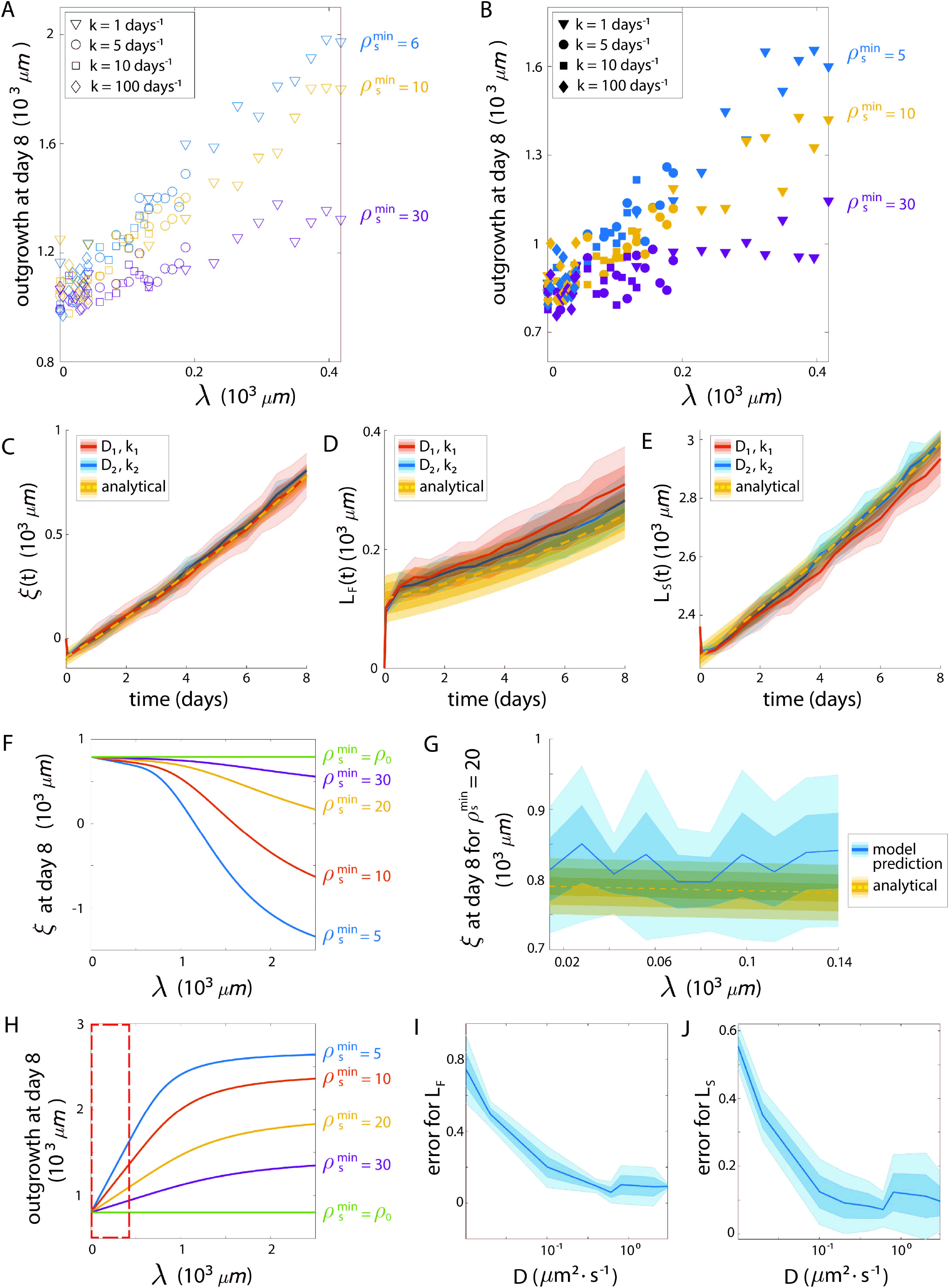
The control of spinal cord regeneration lies in the characteristic length of the signal and the cell-to-signal sensitivity. **(A)** Spinal cord outgrowth predicted by the computational model as function of the characteristic length *λ* of the signal for different values of the signal degradation rate *k* (*k* = 1 day^−1^ (triangle markers), *k* = 5 day^−1^ (round markers), *k* = 10 day^−1^(square markers), *k* = 100 day^−1^ (diamond markers). **(B)** Tissue outgrowth of the computational model assuming Poisson-based cell divisions, same legend as for panel (A). **(C,D,E)** Recruitment limit of the Poisson-based model as a function of time for two sets of parameters (*D, k*) with the same *λ* = 41.74 *μm* (orange and blue) compared with the corresponding analytical solution (yellow dashed line and shadowed areas, each shadowed area corresponds to one ependymal cell diameter) (C). Same representation for the length of the fast-cycling population *L*_*F*_ (*t*) (D) and the slow-cycling population *L*_*S*_(*t*) (E). In (C), (D) and (E), for each set of parameters (*D, k*), means are represented as lines and each shadowed area corresponds to 1 standard deviation out of 5 simulations. **(F,H)** Theoretical predictions of the recruitment limit *ξ*(*t*) (F) and tissue outgrowth (H) as a function of *λ*, for different values of the cell-to-signal sensitivity (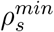). **(G)** Theoretical prediction (yellow) of the recruitment limit *ξ*(*t*) at time *t* = 8 days post amputation as a function of *λ* superimposed with the simulated values using the Poisson-based computational model (blue), for 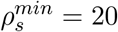 (mean is represented as solid line and each shadowed area corresponds to 1 standard deviation out of 10 simulations). **(I,J)** Error between the theoretical and the numerical values obtained with the Poisson-based computational model of the fast-cycling population length *L*_*F*_ (*t*) (I) and slow-cycling population length *L*_*S*_(*t*) (J) at time *t* = 8 days as function of the diffusion coefficient *D* for fixed value of *λ* = 41.74 *μm*. In (I) and (J) means are represented as lines and each shadowed area corresponds to 1 standard deviation out of 5 simulations.

The cell-based computational model developed in previous sections assumes that the signal recruits cycling cells by reducing G1 and S phase, which accelerates their cell cycle, expanding the spinal cord tissue where the signal operates, as previously reported^7^. Thus, we wondered whether the control of the regenerative response by the characteristic length of the signal and cell-to-signal sensitivity could be an intrinsic feature of the non-Markovian cell cycle dynamics of the cell-based computational model. To explore this hypothesis, we simplified our cell-based computational model by approximating the more correct cell division mechanism with a random (Poisson) process of signal-dependent frequency (see STAR Methods for more details).

Importantly, we observed that the seemingly linear correlation between the regenerative out-growth and the signal characteristic length is preserved (Fig. 4 B). Although the Poisson-based model is definitely less biologically relevant to describe the regenerative response in the axolotl spinal cord, its simplicity makes it ideal to theoretically understand the linear dependence of the tissue outgrowth with the signal characteristic length, where the slope depends on the cell-to-signal sensitivity. In this vein, we developed a theoretical framework of the regenerative growth controlled by a signal whose dynamics is under a reaction-diffusion mechanism.

We formally considered the case of instantaneous signal diffusion and degradation (ideally infinite values of *D* and *k*, corresponding to an instantaneous relaxation of the signal profile towards its steady state). We obtained analytical expressions for the tissue domain occupied by the recruited fast proliferating cells *L*_*f*_ and the domain occupied by non-recruited slow proliferating cells *L*_*s*_ as functions of time (see formal derivation in STAR Methods):

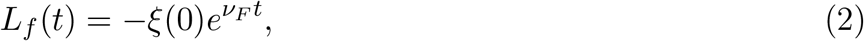

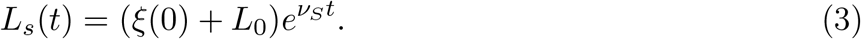

In these expressions, *L*_0_ is the initial length of the spinal cord tissue, immediately after amputation (defined as the distance between the most anterior boundary of the domain and the amputation plane) whereas *ξ*(0) *<* 0 is the position of the recruitment limit at time zero; (*i*.*e*., the spatial point in the AP axis separating the populations of non-recruited and recruited cells (Note that we are using a coordinates system centered in the amputation plane, see Supp. Fig. S3). We found that *ξ*(0) is defined by:

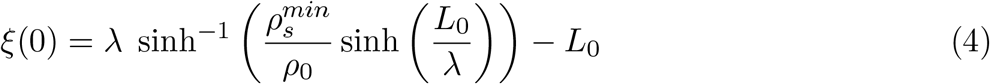

Since the signal lives on the domain occupied by the ependymal cells, we require that *λ < L*_0_. By using Eqs. (2)-(4), we can directly write the theoretical spinal cord outgrowth as a function of time as

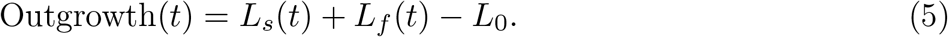

To evaluate the merits of our theory, we decided to test it by comparing the theoretical predictions with simulations of the Poisson-based computational model for large enough values of *D* and *k*. Remarkably, we obtained a very good correspondence between the theoretical values of *ξ*(*t*) (see the general expression of Eq. (4) in STAR Methods), *L*_*s*_(*t*) and *L*_*f*_ (*t*) and their corresponding values computed on the simulations, for all times *t* (Fig. 4 C-E).

This novel theory of spinal cord regeneration allows us to demonstrate how cell recruitment depends on the signal characteristic length as well as the cell to signal sensitivity at a given time (Fig. 4 F). First, by classical asymptotic arguments, one directly obtains that the recruitment limit *ξ*(0) gets closer to 0 immediately after amputation, as the signal characteristic length tends to zero. These results are expected since without diffusion, *D* = 0, the signal is only present on the posterior boundary of the spinal cord and only a small fraction of the population accelerate its cell cycle. Moreover, Eq. (4) reveals how the cell sensitivity to the signal controls the regenerative response. First, notice that 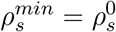 (the density of the signal at the front) leads to *ξ*(0) = 0 (Fig. 4 F). Again, in this case, the cells can only be recruited if the signal density is larger than the one prescribed on the front, i.e., only the cells at the very spinal cord front will accelerate their cell cycles and this leads to the same behavior as a non-diffusive signal (*D* = 0). As 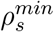 increases, so does *ξ*(0), indicating that higher sensitivity of tissue cells to the signal leads to larger outgrowth (Fig. 4 F). Finally, plotting the theoretical value of the outgrowth given by (5)-(4) as function of *λ* (Fig. 4.H) showed that indeed for small values of *λ* (red box corresponding to the range of *λ* used for our simulations), the outgrowth evolved almost linearly with *λ*, the slope depending on the threshold density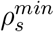.

We further validated the formal derivation by comparing the theoretical dependence of the recruitment limit *ξ*(*t*) on the signal characteristic length with that dependence observed from simulations using the Poisson-based computational model (Fig. 4 G). We obtained a very good agreement between the theory and the simulations for different values of cell to signal sensitivity (Fig. 4 G). It is noteworthy that we were only able to perform simulations for *λ <* 400 *μ*m because of computational time limitations. Indeed, as the characteristic length increases, the numerical time-step required for the simulation decreases, leading to an increase of the overall computation time.

By using our new theory (Eqs. (2) - (5)), we represented the predicted spinal cord outgrowth at a given time (8 days post amputation) as a function of the signal characteristic length, for different cell to signal sensitivities. Remarkably, our results demonstrate that both magnitudes are linearly correlated for characteristic lengths approximately smaller than 400 *μ*m (see the rectangle within Fig. 4 H), with a slope depending on the cell sensitivity to the signal 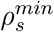, in good agreement with our simulation results (Fig. 4 A, B). However as our theory predicts, the tissue length evolves as a saturated exponential for larger values of *λ*. This saturation effect is due to the finite size of the initial domain for ependymal cells. As the signal invades the domain, it recruit more cells up until reaching the anterior boundary where no more cells can be recruited anymore, saturating the overall evolution of the tissue.

Finally, to investigate the limitations of our theory, we determined the regime within the parameters space in which the theoretical derivation is valid. As the theoretical derivation was made under the regime of instantaneous reaction-diffusion of the signal (*i*.*e*., high diffusion coefficient and degradation rate), we expected that the numerical simulations get closer to the theoretical predictions as the time scale of the reaction-diffusion process dominates the time scale of cell division. To this aim, we arbitrarily fixed the value of the signal characteristic length *λ* = 41.74*μ*m and computed the error, as defined in Eq. (12) in STAR Methods, between the theoretical and simulated tissue domains occupied by the recruited fast proliferating cells *L*_*f*_ and the domain occupied by non-recruited slow proliferating cells *L*_*s*_, for different values of the diffusion coefficient *D*. As expected, we observed that the error decreases as the diffusion coefficient increases (Fig. 4 I, J).

Taken together, the results of this section explain why the regenerative spinal cord outgrowth is approximately linear with the signal characteristic length for a given cell to signal sensitivity. Our theoretical results demonstrate that the regenerative response of the modelled spinal cord in the regime of fast diffusion and reaction is determined by the sensitivity of the cells to the signal, and is not controlled by the diffusion coefficient of the signal and its half-life, but rather by the characteristic length of the signal.

### 2.5 Identification of a signal-dependent transient modulation of cell cycling during axolotl spinal cord regeneration

The previous section showed that the fast reaction and diffusion regime leads to a tissue out-growth controlled by the signal characteristic length and the cell-to-signal sensitivity. This regime is consistent with the regeneration-inducing signal instantaneously reaching its steady state spatial distribution. Nevertheless, the model best-fitting results obtained in section 2.2 and 2.3 indicated a transient evolution of the signal before converging towards the steady state. Indeed, our computational model predicts a signal density, which is higher in the posterior region while propagating anteriorly over time. Thus, the higher posterior values of the signal density recruit ependymal cells by reducing G1 and S phases, transiently leading to more cells in S phase in posterior locations, as previously demonstrated^7^. This is why we sought to validate deeper the model by determining the spatiotemporal distributions of cells in G1/G0 and S/G2 phases from the model simulations and to confront them with the corresponding experimental distributions extracted from regenerating spinal cords of AxFUCCI animals. These transgenic animals allow us to visualize cells in the aforementioned cell cycle phases *in vivo* by using the FUCCI technology^7^. We fixed *λ* = 270*μ* m (*i*.*e*., the best-fitting value of *λ*), by using a signal diffusion coefficient and degradation constant of 0.08 *μ* m^2^sec^−1^ and 0.1 days^−1^, respectively. The computational simulations qualitatively resemble the AxFUCCI regenerating spinal cords (Fig. 5 A, A’). In Fig. 5 B, we show these distributions at different days post-amputation, for different values of (*D, ν*_*d*_) at fixed *λ* = 270*μ* m (*i*.*e*., the best-fitting value of *λ*). As one can observe in Fig. 5 B, the simulations reveal the same behavior as the experimental data: Up until day 2, cells are mostly in the G0/G1 phase, and the proportion of cells in G2/S phase increases at the amputation site, spreading anteriorly from day 3. While we recovered a very good agreement between simulations and experimental data for (*D, ν*_*d*_) = (0.08*μm*^2^.*s*^−1^, 0.1*days*^−1^) (dotted-markers), faster signals failed to recover the correct switch point at days 2 and 3 ((*D, ν*_*d*_) = (8*μm*^2^.*s*^−1^, 10*days*^−1^), round-markers or (*D, ν*_*d*_) = (0.8*μm*^2^.*s*^−1^, 1*days*^−1^), cross-markers). In order to quantify the agreement between the simulations and the data, we computed the relative error between the spatial distributions of cells in G0/G1 (respectively G2/S phase) obtained numerically and with the experiments (see STAR Methods for details on the computation of the error), and plotted this measure as function of time and for different set of parameters in Fig. 5 (C). Indeed, Fig. 5 (C) confirms that the error between the experiments and the simulations is minimal for (*D, ν*_*d*_) = (0.08*μm*^2^.*s*^−1^, 0.1*days*^−1^) (dotted line), compared to faster signals (lines with cross and round markers) which favour faster anterior spreading of the recruitment limit from days 1 to 3. Moreover, consistently with the results shown in previous sections, parameter values leading to different signal characteristic lengths do not recover the correct distributions (Supp. Fig. S4). Therefore, the agreement between the spatiotemporal distribution predicted by the model and the AxFUCCI experiments indicate that the optimal values of the signal parameters are (*D, ν*_*d*_) = (0.08*μm*^2^.*s*^−1^, 0.1*days*^−1^). These results reveal that the signal temporal dynamics is crucial for the kinetics of the distribution of cells in the different phases of their cell cycle. Indeed, even if the signal reaches the same steady state on the spatial scale (with fixed characteristic length *λ*), a faster signal needs less time to establish its steady profile and recruits cells earlier in the regeneration process. To better understand the cell recruitment process, we quantified the number of newly recruited cells by the signal and represented it as function of time (Fig 5 C’). This enabled us to discover two major findings: first, although continuously created at the front of the tissue, we found that the signal only has action (*i*.*e*., actively recruit new cells) during a transient time window. Interestingly, one observes that for each pair of values of (*D, ν*_*d*_), The number of newly recruited cells first increases and then decreases until it reaches 0 in a finite time window, corresponding to the time that the signal only lives on the already recruited cells. Secondly, we observed that the distribution is more picked and happens on shorter timescales when increasing the signal dynamics (*D, ν*_*d*_). We observed a very peaked distribution in the first day post-amputation for (*D, ν*_*d*_) = (8*μm*^2^.*s*^−1^, 10*days*^−1^) while the distribution is spread over 2 days for the slower signal (*D, ν*_*d*_) = (0.8*μm*^2^.*s*^−1^, 1*days*^−1^) and up until 4 days for an even slower signal (*D, ν*_*d*_) = (0.08*μm*^2^.*s*^−1^, 0.1*days*^−1^). Further analysis of this distribution as a function of *D* and *ν*_*d*_ demonstrates that the size of the peak is entirely controlled by the diffusion coefficient *D* (*i*.*e*., increasing diffusion leads to more recruitment, Supp. Fig. S4 B) while the length of the time window is completely determined by *ν*_*d*_ (*i*.*e*., increasing *ν*_*d*_ decreases the time window) (Supp. Fig. S4 D).

**Figure 5:**
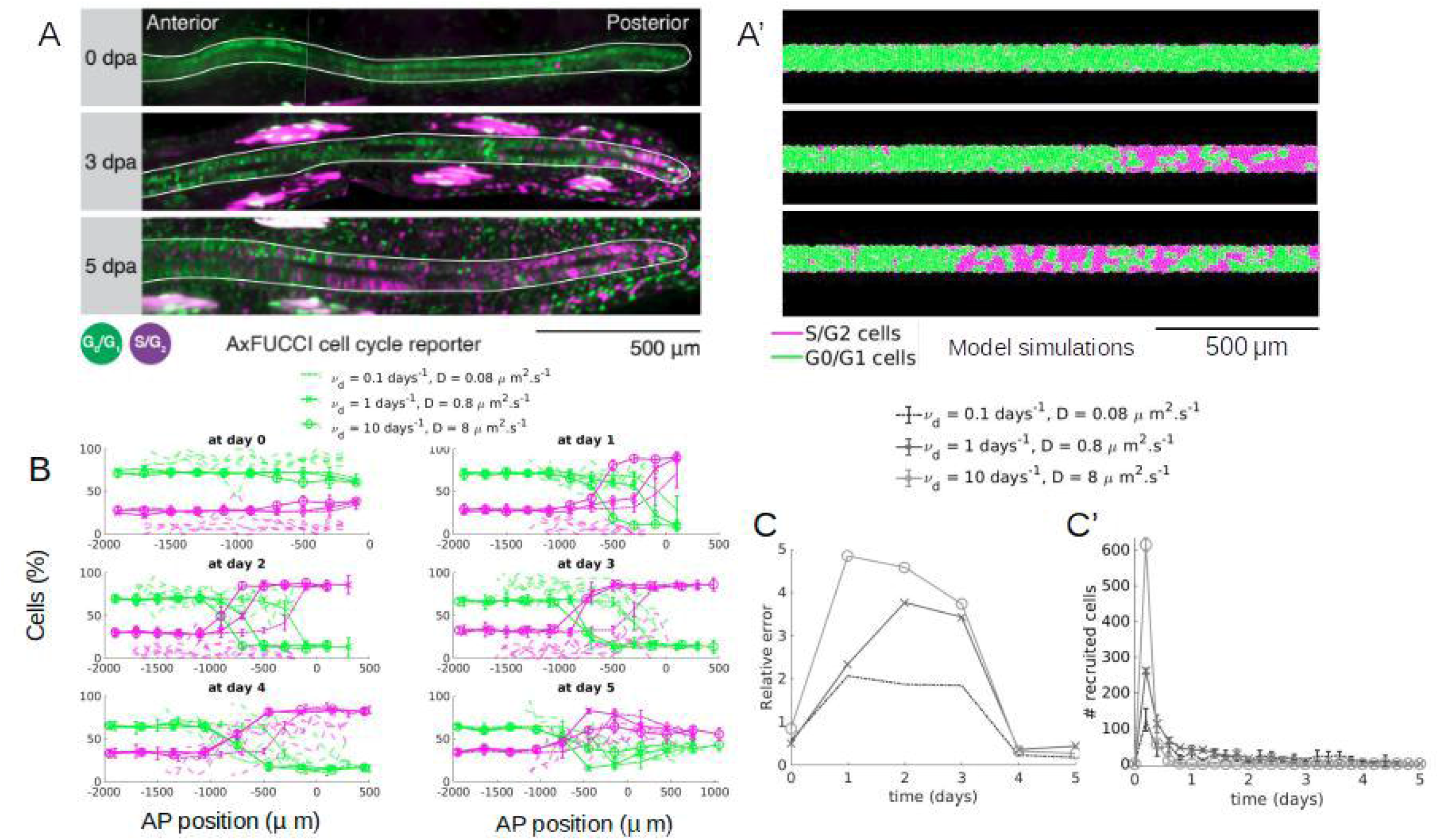
The computational model recapitulates the experimental spatiotemporal distribution of cells in G1/G0 and S/G2 phases obtained in AxFUCCI animals. **(A)** Experimental axolotl spinal cords imaged during regeneration after amputation, using the FUCCI technology, at different days post-amputation (from top to bottom: 0dpa, 3dpa and 5dpa), where cells have been marked according to their position in the cell cycle (green cells in G0/G1 phase, magenta cells in S/G2 phases, for more details see^7^). **(A’)** Same representation with a model simulation for (*ν*_*d*_, *D*) = (0.1*days*^−1^, 0.08*μm*^2^.*s*^−1^). **(B)** Percentage of cells along the AP-axis that are in G0/G1 phase (green curves) and in S/G2 phase (magenta) as function of the AP position, for different times post-amputation (different subplots). Dotted lines are experimental data from^7^, solid lines are simulations for different pairs of (*D, ν*_*d*_) and fixed *λ* = 270*μ* m. For each set of parameters, simulated data are averaged over 10 realisations (solid lines represent the mean, errorbars the standard deviation). **(C)** Relative error (see STAR Methods for details on its computation) between the simulated and experimental cell distributions as function of time post-amputation for the different pairs (*D, ν*_*d*_) (values in the legend) **(C’)** Number of newly recruited cells by the regeneration-inducing signal computed every 0.2 days as function of time post-amputation for the different pairs (*D, ν*_*d*_) (values in the legend). AxFUCCI Images in (A) courtesy of Leo Otsuki and Elly Tanaka and correspond to one of the animals represented in (B)

## 3 Discussion

Severe trauma of the spine leads to irreversible and usually irreparable consequences in most vertebrates. In humans, these traumatic injuries often have dramatic repercussions not only for the patients suffering from those injures but also for their families who take on caring responsibilities for the patients. These overwhelming consequences observed in humans after traumatic injuries in the spine strikingly contrast with the complete structural and functional regeneration of the spinal cord displayed by the axolotl after the most extreme possible injury: tail amputation. Why this salamander is capable of such remarkable regeneration traits while we are not is still elusive^23^.

To address the complexity of the regenerative response observed in the axolotl spinal cord, a first non-spatial mathematical model has been proposed^6^. This first model focused on the kinetics of proliferating and quiescent cells and was encoded in a system of two ordinary non-autonomous differential equations. A subsequent 1D computational model was developed in which cells were simulated as hard segments while the signal was modelled following a phenomenological approach^7^ (For details on the comparison between the model reported in^7^ and the one presented in this study, see Section 2.1 of the Supplementary Information). By combining this simple model with functional experiments using FUCCI technology in axolotls, we have previously shown that spinal cord regeneration in the axolotl is consistent with a process of ependymal cell recruitment triggered by an unknown signal that propagates ≈ 830 *μ*m anteriorly from the injury site during ≈ 85 hours post amputation^7^ However, the signal and the precise mechanism of its propagation within the axolotl spinal cord remain to be elucidated.

During development, numerous tissues are shaped through morphogenetic processes controlled by morphogens or signals whose dynamics are governed by reaction-diffusion processes. As an example, growth regulation of the Drosophila wing imaginal disc is governed by the Decapentaplegic (Dpp) morphogen gradient, in turn ruled by a reaction-diffusion mechanism^24^. In vertebrates, a reaction-diffusion system explains the dynamics of the transforming growth factor–*β* superfamily signals Nodal and Lefty during zebrafish embryogenesis^25^. In a chick embryo, a reaction-diffusion mechanism was proposed to explain the antagonistic effect of BMP2 and BMP7 on the feather patterning^26^. These and other examples show that the reaction–diffusion models constitute effective and accurate mathematical constructs guiding mathematical approaches in development^27^ and possibly in regeneration^12^. Thus, it is conceivable that the particular spatiotemporal distribution of the signal responsible for the regenerative response observed in the axolotl spinal cord might reflect a reaction-diffusion mechanism at work. This is why, in this theoretical study, we decided to test this hypothesis.

To that aim, we have developed a 2D multi-scale hybrid computational model featuring ependymal cells which interact with a signal continuously produced at the posterior tip of the spinal cord, which diffuses within the tissue while being consumed with a given degradation rate. When the local signal density overcomes a certain threshold, the ependymal cells in contact with the signal accelerate their cell cycle, increasing the overall tissue growth speed (Fig. 1). After validating both the cellular and the signalling scales individually (Supplementary sections 2.1 and 2.2), we tested the model by fitting the previously reported recruitment limit curve^6^. Interestingly, the model is able to recover the spinal cord outgrowth experimentally observed^6^ (Fig. 2 F, G, Fig. 3 B, C, Supp. Fig. S1 B), and comparison with AxFUCCI experimental data allowed us to have a first estimate of the biophysical parameters governing the hypothetical signal responsible for the regenerative process observed in the axolotl spinal cord. The predicted signal diffusion coefficient and degradation rate are ≈ 0.1*μ*m^2^sec^−1^ and 0.1*days*^−1^, respectively. While there are no measurements of diffusivity or stability of morphogenetic signals in the axolotl during regeneration, these biophysical parameters were determined for a variety of morphogens in zebrafish embryos *in vivo*. As an example, the diffusion coefficients of Cyclops, Squint, Lefty1 and Lefty2 were estimated in (*μ*m^2^sec^−1^) 0.7 *±* 0.2, 3.2 *±* 0.5, 11.1 *±* 0.6, and 18.9 *±* 3.0, respectively, while their half-lives ranged from (min) 95 to 218 in the blastula stage during zebrafish embryogenesis^25^. Interestingly, in the same animal model, BMP2b and Chordin have clearance rate constants of (10^−5^*s*^−1^) 8.9 *±* 0.1 and 9.6 *±* 0.3, corresponding to half-lives of (min) 130 and 120, and effective diffusion coefficients of (*μ*m^2^sec^−1^) 2–3 and 6–7, respectively^28^. Hence, the diffusion coefficient and half-life of the hypothetical signal predicted by our model are not very different from those observed in Cyclops and BMP gradients determined in the zebrafish embryo. These differences could be attributed to the differences in animals, stage and anatomical orientation (anterior-posterior axis in the axolotl spinal cord versus dorsal-ventral axis in the zebrafish embryo) or they could of course reflect different signals.

Importantly, our computational model results indicate that the regenerative growth response scales with the characteristic length of the signal, where the slope depends on the ependymal cell-to-signal sensitivity (Fig. 4 A), result that was also recapitulated when the proliferative response of the ependymal cells was simulated with a naïve Poisson-based proliferation model (Fig. 4 B). To explain this result, we developed a theory that allowed us to unveil a profound feature of the regenerative response displayed by the computational model. Our rigorous mathematical demonstration reveals that the spinal cord outgrowth emerging during regeneration in the axolotl can be controlled by the characteristic length of the signal rather than by its individual diffusion coefficient or half-life. This study predicts that the signal governing the regenerating spinal cord in the axolotl has a characteristic length of about 270 *μ*m. Interestingly, gradients of BMP and Chordin display characteristic lengths of 168 and 260 *μ*m, respectively (calculated from^28^), suggesting that the our regeneration-inducing signals could operate within the BMP pathway.

Our model predicts that a reduction in the diffusion coefficient would result in a shortened spatial expansion of recruited cells, reflected in a correspondingly shortened posterior domain dominated by S-phase cells, leading to a reduced regenerative response. Although the identity of the regeneration-inducing signal remains to be elucidated, the diffusion coefficient suggests a molecular size similar to that of BMP or Chordin. Interestingly, a difference in diffusion coefficient has been reported for BMP using FRAP and FCS, which has been attributed to differential binding to extracellular molecules^28^, as has been proposed for other developmental signals such as Nodal and FGF^25,29^. Similarly, according to our model, reducing the half-life of the regeneration-inducing signal would be the expected result of upregulating the antagonist of the regeneration-inducing signal. Indeed, BMP gradients are modulated by a network of extracellular regulators in both vertebrates and invertebrates^30,31^. In the context of our model, modulation of the diffusion coefficient or half-life could lead to recruitment and thus to a regenerative response that would ultimately be indistinguishable from up- or down-regulation of the signal itself.

The existence of a transient time window of newly recruited ependymal cells observed with the model simulations (Fig. 5 D) is reminiscent of the kinetics of AxMLP protein expression measured in the axolotl upon tail amputation^32^. Indeed, the AxMLP protein levels is upregulated with a peak of expression at 12 to 24 h post-amputation, returning to basal levels at 2 days post-amputation^32^.

Noteworthy, the reaction-diffusion mechanism under consideration here may be an effective mechanism arising from a process which does not necessarily involve morphogen gradients. Indeed, there is a growing appreciation that morphogenetic processes can be moulded by physical cues^33^. A clear example is the fluid-to-solid jamming transition observed during posterior axis elongation in zebrafish embryos^34^. In this system, there is a transition of rigidity from anterior to posterior tissues where the fluid-like posterior tissues remodel before their maturation. A similar process takes place during development of the zebrafish blastoderm where the rigidity transitions from a network of high to low intercellular adhesions^35^. Hence, the reaction-diffusion mechanism governing our hypothetical signal responsible for spinal cord regeneration after amputation could correspond to a mechanical process. Interestingly, a mechanical reaction-diffusion mechanism elegantly explains wound-induced regeneration of hair follicles in mice^36^. In fact, the idea was already anticipated in the seminal study of George Oster, James D. Murray and Albert

K. Harris^37^ that showed pattern formation emerging from cell motility and rigidity perturbations, very much like reaction-diffusion instabilities predicted by the Gierer-Meinhardt^38^ and the Turing models^39^.

### 3.1 Limitations of this study

It is noteworthy that this simple 2D model can be reduced to a 1D model due to the choice of boundary conditions (periodic in the y-direction) and spatially homogeneous rules for cell divisions. Future perspectives considering different domain geometries will take into account the variation of the spinal cord diameter along the AP axis and study its impact on tissue outgrowth. The extension to 3D will allow more realistic mechanical interactions to be taken into account, reflecting both cell-cell stresses and intracellular pressures. The 2D cell-based computational model developed here focuses on the early regenerative response of the spinal cord in the axolotl. A control mechanism via a negative feedback process that effectively turns down proliferation, and thus regeneration, must be present within the first few weeks after amputation. Indeed, asymmetric cell divisions of ependymal cells are observed during the second week of regeneration^5^. Since the dynamics of newly generated neurons occur along the radial coordinate of the ependymal tube (perpendicular to the AP axis), a 3D model would be advisable to study the regenerative response of the spinal cord in the long term. A possible strategy to circumvent the resulting complexity of the spinal cord during this later regulatory phase of regeneration could involve mapping the cell-based computational models with a macroscopic (continuous) description, as those used in^40,41,42^. Additionally, the theory demonstrating that the regenerative growth scales with the characteristic length of the signal relies on the fast diffusion and reaction approximation. Future studies should address whether this scaling extends to other reaction-diffusion regimes.

This study shows that spinal cord regeneration in the axolotl can be orchestrated by a hypothetical signal operating under a reaction-diffusion scheme under the control of its characteristic length and the sensitivity of the ependymal cells to the signal. Further investigation is required to unveil the nature of the signal to then explore how conserved and thus general this mechanism of regeneration might be.

## Supporting information

Supplementary section

## 4 Acknowledgements

D.P. was supported by Sorbonne Alliance University with an Emergence project MATHREGEN, grant no. S29-05Z101 and by Agence Nationale de la Recherche (ANR) under the project grant number ANR-22-CE45-0024-01. OC was funded by grants PICT-2017-2307 and PICT-2019-03828) from the Agencia Nacional de Promoción Científica y Tecnológica of Argentina. The authors warmly thank Leo Otsuki and Elly Tanaka for the experimental images of the AxFUCCI animals depicted in Fig. 5 (A). The authors sincerely thank Alberto Ceccarelli, Leo Otsuki, Sophie Hecht and Fabian Rost for their careful reading of the manuscript and their excellent suggestions, as well as Elly Tanaka for incessant and stimulating discussions.

## 5 Author contributions

V.C. developed the computational models, performed and analyzed the simulations and wrote the first draft of the manuscript. D.P. developed the mathematical theory. D.P and O.C. conceived the project, supervised V.C., secured funding, analyzed the results and wrote the final study.

## 7 Main Figure titles and legends

## 8 STAR Methods

### RESOURCE AVAILABILITY

#### Lead Contact

Further information and requests for resources should be directed to Osvaldo Chara.

#### Materials availability

This study did not generate new materials.

#### Data and Code availability

- Data: This is a computational study. No new experimental data has been generated.
- Code: The computational model in this study was implemented in Fortran 90 while visualization of simulations and analyses were executed with MATLAB. All original codes for simulations and post-processing have been deposited at https://doi.org/10.5281/zenodo.8111099
- Any additional information required to reanalyze the data reported in this work paper is available from the Lead Contact upon request.

### METHOD DETAILS

#### A mathematical 2D hybrid model for spinal cord repair after injury

The 2D mathematical hybrid model features a number *N* (*t*) of ependymal cells represented as individual 2D hard-disks of centers *X*_*i*_(*t*) and uniform and fixed radii *R* for *i* = 1 … *N* (*t*), and a signal represented as a continuous concentration field *ρ*_*s*_(**z**, *t*) depending on space and time. The ependymal cells live in a 2D domain Ω(*t*) = [−*L*_0_, *L*(*t*) −*L*_0_] *×* [*y*_min_, *y*_max_], where *L*(*t*) indicates the size of the tissue at time *t* and *L*_0_ = *L*(0), aiming at modelling the spinal cord apical surface as the surface of a cylinder (see Fig. 1A). Note that the coordinate system is centered in the amputation plane. We equip this domain with periodic boundary conditions in the <u>y</u>-direction (representing the spinal cord circumference), Dirichlet boundary condition on the left (anterior part of the tissue supposed to be fixed) and free boundary on the right (posterior/amputated part of the tissue that can grow). The signal is supposed to be produced at the posterior tip of the tissue (the free boundary on the right in Fig. 1A), degrade with constant degradation rate *k ∈* R^+^ and diffuse in the domain occupied by the ependymal cells Ω(*t*).

##### Reaction-diffusion equation for the signal

These simple assumptions lead to the following reaction-diffusion equation for the signal density *ρ*_*s*_(**z**, *t*):

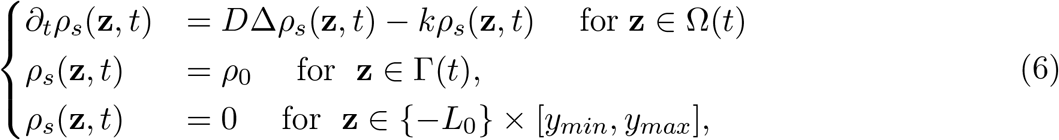

where Γ(*t*) = *{L*(*t*) − *L*_0_*} ×* [*y*_min_, *y*_max_] is the front of ependymal cells. The tissue length *L*(*t*) is determined by the average of the <u>x</u>-coordinates of the front cells (see definition of front cells in Section “Identification of front cells”).

##### Ependymal cells recruitment and division

Cells surrounded by a signal concentration above a certain threshold 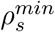 are supposed to be instantaneously recruited. We sketch cell recruitment as function of the signal in Fig. 1B, where we show the signal concentration as function of the AP-axis position, where the amputation plane is indicated with a vertical dashed line. Cells are supposed to cycle with given cell cycle lengths drawn from lognormal distributions *Log*(*μ, σ*) which parameters depend upon their status (long cell cycle length with mean *μ* = 340h and standard deviation *σ* = 32 h for non-recruited cells, small cell cycle length with mean *μ* = 119h and standard deviation *σ* = 10h for recruited cells), and divide when reaching the end of their cell cycle. We assume that, when recruited, an ependymal cell shortens its cell cycle by partial skipping of the *G*_1_ phase and proportional mapping between long and short *S* phases (see^7^ for more details).

Note that the ependymal cells recruited by the signal reduce G1 and S phases, effectively shortening the cell cycle and inducing partial synchronisation. Since the modelled signal is generated in the posterior tip and propagates anteriorly by the reaction-diffusion mechanism, cells progressively acquire partial synchronisation from posterior to anterior.

We consider that cell growth is faster than any other mechanisms and therefore upon division, an ependymal cell of radius *R* produces instantaneously a new daughter cell of the same radius positioned randomly in a disc centered around its center and of radius 2*R*. When the ependymal cell *i* of center *X*_*i*_ has reached the end of its cycle, it generates two ependymal cells *j* and *k*: cell *j* is placed at *X*_*j*_ = *X*_*i*_, while cell *k* is placed at a random distance in (0, 2*R*) from the cell *j* and with an angle *θ* chosen randomly in (−*θ*^*\**^, *θ*^*\**^). We checked that the choice of *θ*^*\**^ does not impact the outcome of the process (data not shown), and arbitrarily set 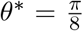 in this paper. We suppose that cell recruitment is irreversible and inherited from mother to the daughter cells, <u>i.e.</u>, once recruited, a cell and all its progeny will stay recruited. We suppose that ependymal cells behave as hard discs, <u>i.e.</u>, they move instantaneously to avoid the overlapping induced by the division of cells (see next section for the numerical details).

Therefore in this model, tissue growth is induced by ependymal cell proliferation, increasing the internal pressure of the system that expands to avoid overcrowding. The cell proliferation is itself controlled by the presence of a signal that diffuses from the front of tissue cells towards the anterior side of the AP-axis, and recruits irreversibly new cells as it diffuses in the tissue. As tissue grows, the signal profile (living on a growing domain) is shifted to the posterior axis, until it only lives on already recruited cells.

We give in Supplementary table S1 the model parameters we used for our simulations. The cell cycle lengths *μ*_*S*_ and *μ*_*F*_ for slow and fast cycling cells (respectively) are taken from measurements in^5^. As we don’t have access to direct measurements for our signal (i.e for the diffusion coefficient *D* and degradation rate *k* as well as cell sensitivity to the signal 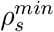), we consider them as free parameters and explore a broaden range of values. Note that the values we consider are still in the range of realistic experimental measurements for morphogens diffusion and degradation rates extracted in different types of tissues^43^.

#### Numerical methods

A major difficulty in the simulation of systems of large number of particles is the high computational cost, typically quadratic in the number of particles. In order to reduce the computational cost of our simulations, we used localization techniques to compute the interactions and implemented the scheme in Fortran90, offering high efficiency, computational precision, vast libraries of matrix, physics and engineering functions and parallelization tools enabling to perform high precision simulations.

##### Initial conditions

We initialize the model with *N* ependymal cells in non-overlapping configuration in the domain [0, *L*_0_]*×*[*y*_*min*_, *y*_*max*_]. This is achieved by throwing randomly *N*_1_ *> N* cells in [0, *L*_0_]*×*[*y*_*min*_, *y*_*max*_], letting the system reach the steady-state of Eq. (9) (non-overlapping configuration), and removing all cells which *x*-position is above *L*_0_. We suppose that initially, all ependymal cells are proliferating slowly and each cell cycle length *T*_*i*_(0) is drawn from a lognormal distribution with mean *μ* = 340*h* and standard deviation *σ* = 32*h* (see supplementary table S1 for more details on the parameter values). Each cell has initial age *C*_*i*_(0) drawn randomly from an exponential distribution given by 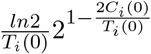.

After this initialization step for the ependymal cells, we initialize the signal by distributing randomly *N*_*S*_ signalling particles close to the amputation plane, more precisely in the domain [*L*_0_ − 2*R, L*_0_] *×* [*y*_*min*_, *y*_*max*_] where *R* is the radius of an ependymal cell. The number of signalling particles *N*_*S*_ is chosen to account for the Dirichlet condition *ρ*_*S*_ = *ρ*_0_ at the boundary, which gives:

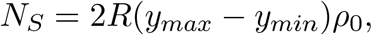

where *ρ*_0_ is the density of the signal imposed at the front (see supplementary table S1 for more details on the parameter values).

##### Reaction-diffusion of the signal

In order to solve the reaction-diffusion equation (6), we use a splitting method. In the first splitting step, we use the Smoothed Particle Hydrodynamics (SPH) method to solve for the diffusion term (first term on the right hand side of Eq. (6)), and in the second step, we solve for the reaction term (second term on the right hand side of Eq. (6)). The choice of a SPH method is motivated by the initially zero signal concentration. Regions of zero concentration are difficult to handle with classical grid methods such as finite volume methods, since it often leads to the appearance of negative values breaking down the simulations. The SPH method is not subject to this risk and is the method of choice for problems involving jets or injections of gases in vacuum. This method has been extensively studied and its accuracy has been practically assessed^44,20,45^. For the first step, We discretize the signal by means of *N*_*S*_ particles, the so-called signalling particles, of mass *m*_*j*_ placed at *Z*_*j*_(*t*) *∈* Ω(*t*), as done by^46^. The density of the signalling particles, governed by diffusion, evolves in time as follows:

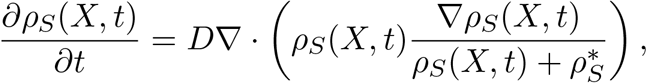

Where we have introduced the constant parameter 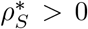 prevent any singularity in case of *ρ*_*S*_ = 0. Switching to a Lagrangian description of the fluid and following the individual particles through which the continuum field has been discretized, we describe the time evolution of the position *Z*_*j*_ of particle *j* as follows:

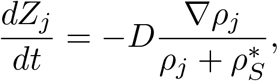

where *ρ*_*j*_ = *ρ*_*S*_(*Z*_*j*_) and *∇ρ*_*j*_ = *∇ρ*_*S*_(*Z*_*j*_) are computed using the Smoothed-Particle Hydrodynamics (SPH) method^47^,^48^ as follows

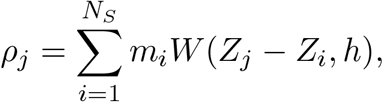

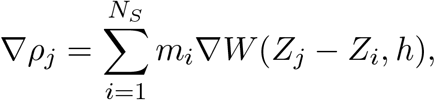

where *W* is the ‘Poly6’ kernel proposed by^29^, adapted to a 2D domain, with support radius *h*, which is defined as follows

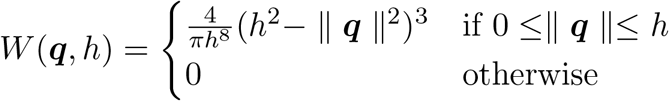

and whose gradient is

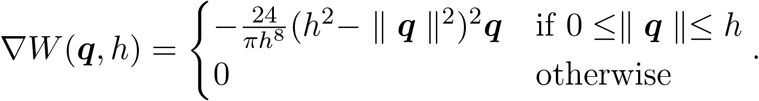

Finally, considering an explicit time discretization, we define *t*^*n*^ = Δ*t*^1^ + … + Δ*t*^*n*^. Then, the position of particle *j* at time *t*^*n*+1^ can be computed as follows:

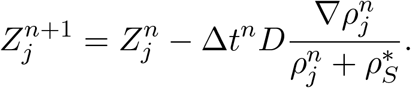

The time step Δ*t*^*n*^ must be chosen such that

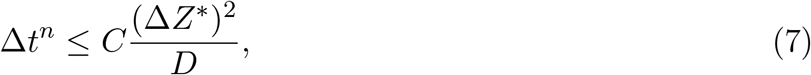

with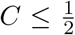, to ensures that, during the time step Δ*t*^*n*^ each particle *j* can move at most Δ*Z*^*\**^, where Δ*Z*^*\**^ was chosen equal to the diameter of an ependymal cell, i.e. Δ*Z*^*\**^ = 2*R*.

We assume that the signalling particles are removed following a Poisson process of frequency *k*, i.e. the probability of a signalling particle to disappear in the time interval (*t, t* + *dt*) is

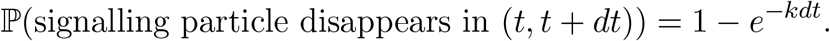

In order to obtain a good approximation of Poisson process, the time step Δ*t*^*n*^ must be chosen such that

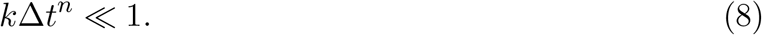

Because of the two conditions introduced in Eqs (7) and (8), the time step Δ*t*^*n*^ is chosen as follows:

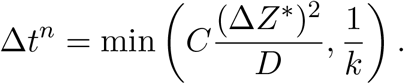

##### Ependymal cells interaction and motion

As explained in the modelling section, cell motion is supposed to be an instantaneous mechanism, i.e at all times, cells are supposed to be in non-overlapping configuration. Numerically, this is treated according to the following scheme: Given a configuration 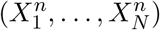 of *N* non-overlapping cells at time *t*^*n*^ undergoing *M* divisions between *t*^*n*^ and *t*^*n*+1^ = *t*^*n*^ +Δ*t* positioned at 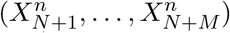 (inducing some overlapping), the configuration of the *N* + *M* cells at time *t*^*n*+1^ is the steady state of

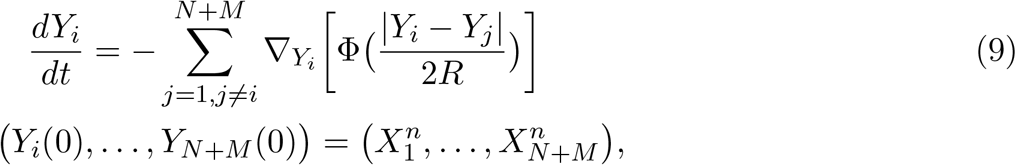

where the repulsion interaction potential Φ is defined as follows:

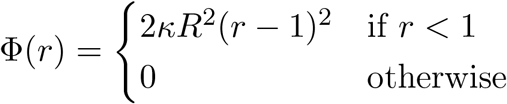

with *i, j* = 1, …, *N* (*t*), where *κ ∈* ℝ^+^ is the repulsion intensity. Note that taking *X*^*N*+1^ to be the steady-state of the dynamics (9) amounts to consider that cells behave as hard spheres, i.e they are in non-overlapping configurations at all times.

Numerically and in between each time *t*^*n*^ and *t*^*n*+1^, we solve (9) using a classical explicit Euler scheme. In order to identify the steady-state, we fix a tolerance *ε >* 0 and we let the repulsion dynamics run until the following stopping criterion is satisfied:

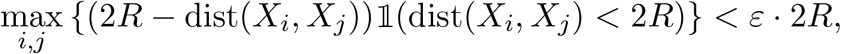

where dist(*X*_*i*_, *X*_*j*_) is the standard Euclidean distance, *R* is the radius of an ependymal cell and 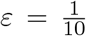. Using this criterion, a non-overlapping configuration is reached when the maximum overlapping between two ependymal cells is less than the 10% of the cell diameter.

##### Ependymal cell division in the Poisson-based model

As described in the results section of the main text, we aimed to test whether the results of our complete model could be an intrinsic feature of the non-Markovian cell cycle dynamics. To explore this hypothesis, we consider a simplified version of our cell-based computational model by approximating the more correct cell division mechanism by a Poisson process of given frequency *ν*_*S*_ (for slow cycling cells) or *ν*_*F*_ (fast cycling cells) with *ν*_*F*_ *< ν*_*S*_. Numerically, this is treated by defining the probability of a fast-cycling (FC) cell to divide between times *t*^*n*^ and *t*^*n*^ + Δ*t* as:

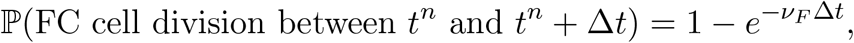

and the same expression holds with *ν*_*S*_ for a slow cycling cell.

#### Theoretical study of the model in the regime of fast reaction-diffusion

In this section, we give the details of the derivation of the model in the regime of fast diffusion and degradation of the signal. We first simplify the model and consider that cell division follows a Poisson process of given frequency *ν*_*S*_ (for slow cycling cells) or *ν*_*F*_ (fast cycling cells) with *ν*_*F*_ *< ν*_*S*_ (see previous section ‘Numerical methods’ for details on the numerical implementation of the Poisson-based computational model). We further simplify the model by considering the 1D case, but the arguments hold in 2D. For the sake of simplicity, we consider shifted x-coordinates and work with the new spatial variable *x* = *z* + *L*_0_ *∈* [0, *L*(*t*)] such that the origin of the new coordinate system is given by the left boundary of the tissue.

We consider the regime of fast diffusion and degradation of the signal, while all the other mechanisms stay of order 1. To this aim, we introduce a small *ϵ* ≪ 1 and set 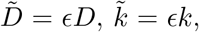, 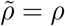 where 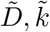 and 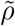 are of order 1 and all the other variables stay of order 1. We get

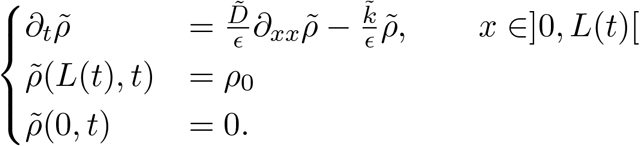

while the dynamics for the cells stay unchanged. We call *L*(*t*) the size of the tissue. Therefore, we obtain that formally as *ϵ* → 0 (omitting the tildes for the sake of clarity)

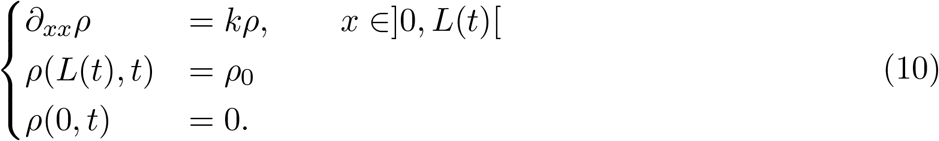

Therefore, at each time *t, ρ* is the steady state of the reaction diffusion equation with Dirichlet boundary conditions on the domain [0, *L*(*t*)]. Solving for *ρ*, we get

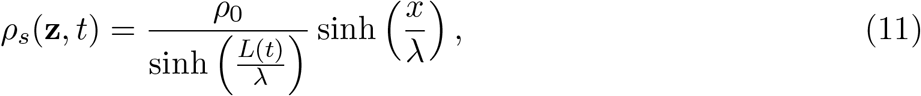

where 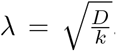. This expression allows us to extract the *x* position denoted *x*_0_(*t*) at which 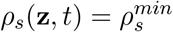:

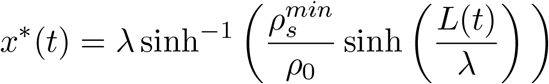

Therefore, in this regime, the signal instantaneously relaxes to its steady-state profile, depending on the tissue length *L*(*t*). At initial time, where *L*_0_ is the tissue length, all the ependymal cells surrounded by a chemical concentration above 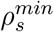 will be instantaneously recruited. Therefore, the initial condition will be composed of slow cycling cells on [0, *x*_0_[and fast-cycling cells on [*x*_0_, *L*_0_]. As both populations divide, this shifts the boundary of the tissue *L*(*t*), instantaneously leading to a new signal profile defined by (11). We now denote by *L*_*S*_(*t*) the length of the slow-cycling population (living on [0, *L*_*S*_(*t*)]), and by *L*_*F*_ (*t*) the length of the fast-cycling population (living on]*L*_*S*_(*t*), *L*_*S*_(*t*) + *L*_*F*_ (*t*)]).

We now claim the following

##### Proposition 1.

*If the initial condition is such that*

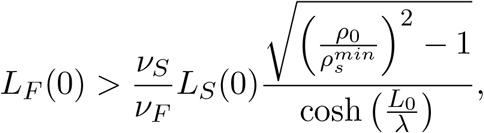

*Then*

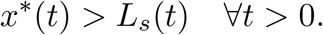

This proposition indicates that the position *x*^*\**^(*t*) after which cells might be recruited lives on already recruited cells for all times *t >* 0. We now proceed with the proof

*Proof*. As cell division is the only phenomenon leading to tissue expansion, the tissue outgrowth is given by

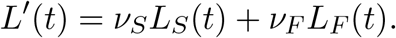

We therefore have to prove that *x*^*\**^(*t*) moves faster than the expansion of the slow-cycling population given by *ν*_*S*_*L*_*s*_(*t*). We compute:

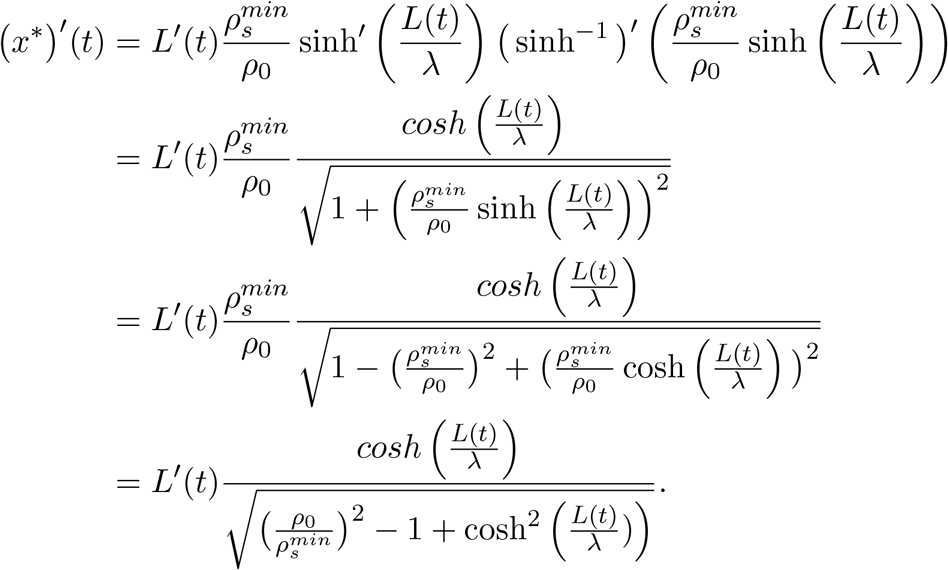

Therefore, the condition *x*^*′*^(*t*) *> ν*_*S*_*L*_*S*_(*t*) amounts to

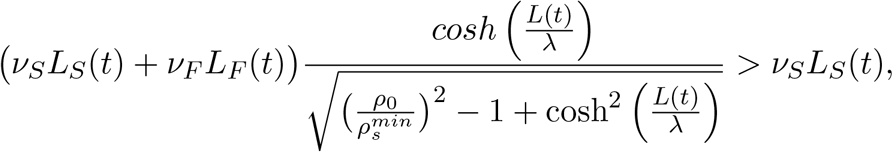

leading to

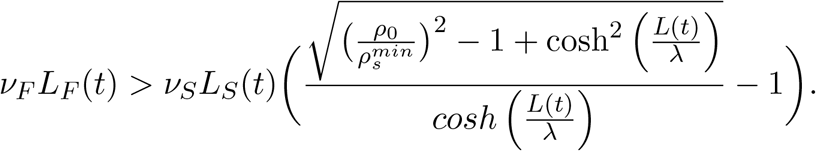

Therefore, as the fast-cycling population expands faster than the slow-cycling one (*ν*_*F*_ *> ν*_*S*_), we need:

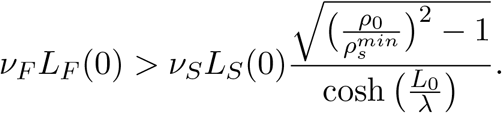

Under the initial condition given by proposition 1, we therefore showed that only the initial cells will be recruited as the recruitment zone lives on already-recruited cells for all times *t >* 0. Therefore, we can compute the evolution of the population of fast- and slow-cycling cells as:

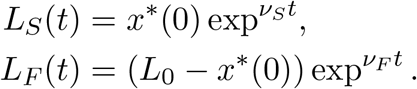

Note that in the original variables, *ξ*(*t*) = *x*^*\**^(*t*) − *L*_0_.

### QUANTIFICATION AND STATISTICAL ANALYSIS

#### Model fitting to the experimental recruitment limit curve

In this section we give details on the procedure we use to fit the model to the previously reported experimental switchpoint^6^.

We denote by 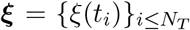 the simulated recruitment limit and with 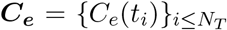 the experimental recruitment limit curve^6^. Then, we introduce 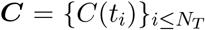 with *C*(*t*_*i*_) = |*ξ*(*t*_*i*_) − *C*_*e*_(*t*_*i*_)| and we define the error between the two curves as follows:

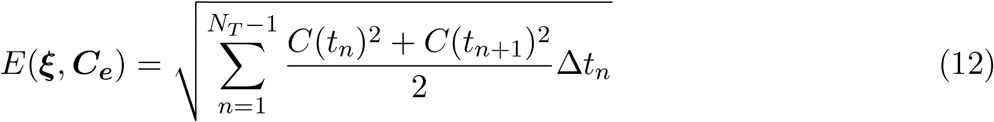

where Δ*t*_*n*_ = *t*_*n*+1_ − *t*_*n*_.

For each combination of values of the signal diffusion coefficient *D* and degradation rate *k* we compute the error, defined in Eq. (13), between the recruitment limit predicted by our model and the one experimentally measured by^6^. We minimize the error by fixing a tolerance *ε* and identifying, among the combinations of values used to perform the simulations, the pairs of *D* and *k* such that *E*(***ξ, C***_***e***_) *< ε* where ***ξ*** is the simulated curve and ***C***_***e***_ is the experimental curve^6^ and *ε* = 0.5 is chosen arbitrarily. This procedure allows to fit the recruitment limit experimentally measured by^6^ with the model-predicted recruitment limit and to identify the combinations of *D* and *k* which best recover the experimental recruitment limit (Fig. 2 D).

### 8.1 Error between simulated and analytical curve

#### General definition

In this section we give the definition of the error between a simulated and an analytical curve. We denote with 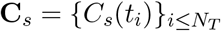 the simulated curve and with 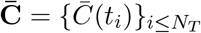 the analytical curve. Then, as done in the previous section, we introduce 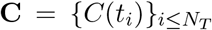 with 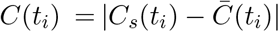 and we define the error between the two curves as follows:

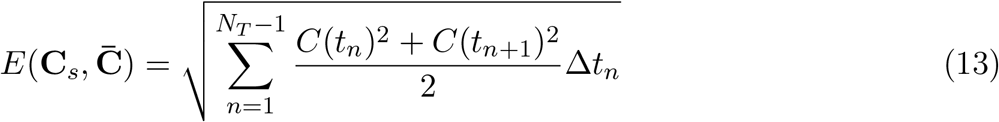

where Δ*t*_*n*_ = *t*_*n*+1_ − *t*_*n*_.

#### Relative error between the simulations and AxFucci experimental data (Fig5 of the main text and FigS5)

Let 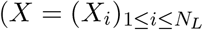 being a vector of points indicating the spatial position along the AP-axis and 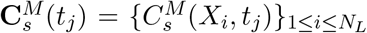 (resp. 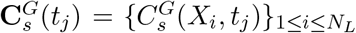) the spatial distributions of cells in S/G2 phase (resp G0/G1 phase) obtained with a simulation at time *t*_*j*_. We denote by 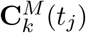 (resp.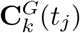) the experimental spatial distributions of cells in S/G2 phase (resp G0/G1 phase) of animal number *k*, interpolated on the points *X*_*i*_. Then, the relative error *E*(*t*_*j*_) between the simulation and experiments is defined by:

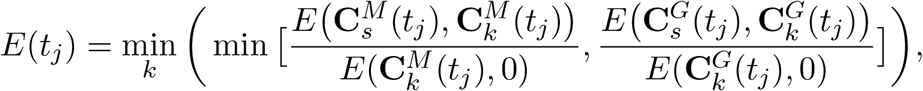

where the minimum is taken over all experiments.

### Identification of front cells

We identify the cell *i* as a front cell if there is no cell *j* such that

- the cell *j* is placed to the right of cell *i*, i.e. *x*_*j*_ *> x*_*i*_,
- *y*_*j*_ *∈* (*y*_*i*_ − *R, y*_*i*_ + *R*), with *R* the cell radius.

For example, in Supp. Fig. S1.A, the ependymal cell *i* (cyan disc) is a front cell because no other ependymal cell (red disc) placed to its right has its center placed in (*y*_*i*_ − *R, y*_*i*_ + *R*). Instead, in Supp. Fig. S1.B, the ependymal cell *i* (blue disc) is not a front cell since the there is a ependymal cell *j* (yellow disc) such that *x*_*j*_ *> x*_*i*_ and *y*_*j*_ *∈* (*y*_*i*_ − *R, y*_*i*_ + *R*).

## 9 Legends of Supplementary Movies

**Supplementary Movie 1: Simulation of the modelled regenerating spinal cord after tail amputation**. Related to Figure 2 and Section 2. Signal diffusion coefficient *D* = 1 *μm*^2^ *· s*^−1^, degradation rate *k* = 1 days^−1^ and 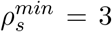. Ependymal cells are represented as discs depicted in three colors: blue, if they are non recruited (slow-cycling) and not exposed to the signal; white, if they are non recruited (slow-cycling) even if exposed to the signal; red, if they are recruited (fast cycling cells). Smaller cyan discs represent the signalling particles. The black dashed line represents the amputation plane.

**Supplementary Movie 2: The evolution of spinal cord outgrowth is controlled by the interplay between the diffusion coefficient** *D* **and the degradation constant** *k*. Related to Figure 2 and Section 2. Simulations of the modelled regenerating spinal cord after tail amputation: large diffusion (*D* = 2.02 *μm*^2^ *· s*^−1^) and large degradation (*k* = 100 days^−1^) of the signal (top), large diffusion (*D* = 2.02 *μm*^2^ *· s*^−1^) and small degradation (*k* = 1 days^−1^) of the signal (middle), small diffusion (*D* = 0.02 *μm*^2^ *· s*^−1^) and small degradation (*k* = 1 days^−1^) of the signal (bottom). The cell-to-signal sensitivity is fixed in 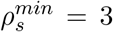. The black dashed line represents the amputation plane. Ependymal cells are represented as discs depicted in three colors: blue, if they are non recruited (slow-cycling) and not exposed to the signal; white, if they are non recruited (slow-cycling) even if exposed to the signal; red, if they are recruited (fast cycling cells). Smaller cyan discs represent the signalling particles.

**Supplementary Movie 3: Tissue regeneration can be modulated by the cell sensitivity to the signal**. Related to Figure 3 and Section 2. Simulations of the regenerating spinal cord after tail amputation. Signal diffusion coefficient *D* = 1 *μm*^2^ *· s*^−1^ and degradation rate *k* = 1 days^−1^. Ependymal cells are represented as discs depicted in three colors: blue, if they are non recruited (slow-cycling) and not exposed to the signal; white, if they are non recruited (slow-cycling) even if exposed to the signal; red, if they are recruited (fast cycling cells). Smaller cyan discs represent the signalling particles. At the top, for large sensitivity to the signal (small 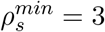), a large zone of high proliferative cells is observed. At the bottom, for small sensitivity to the signal (large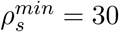), only cells close to the front are activated.

