## Supplementary section for "How a reaction-diffusion signal can control spinal cord regeneration in axolotls: A modelling study"

### 1 Supplementary figures and tables

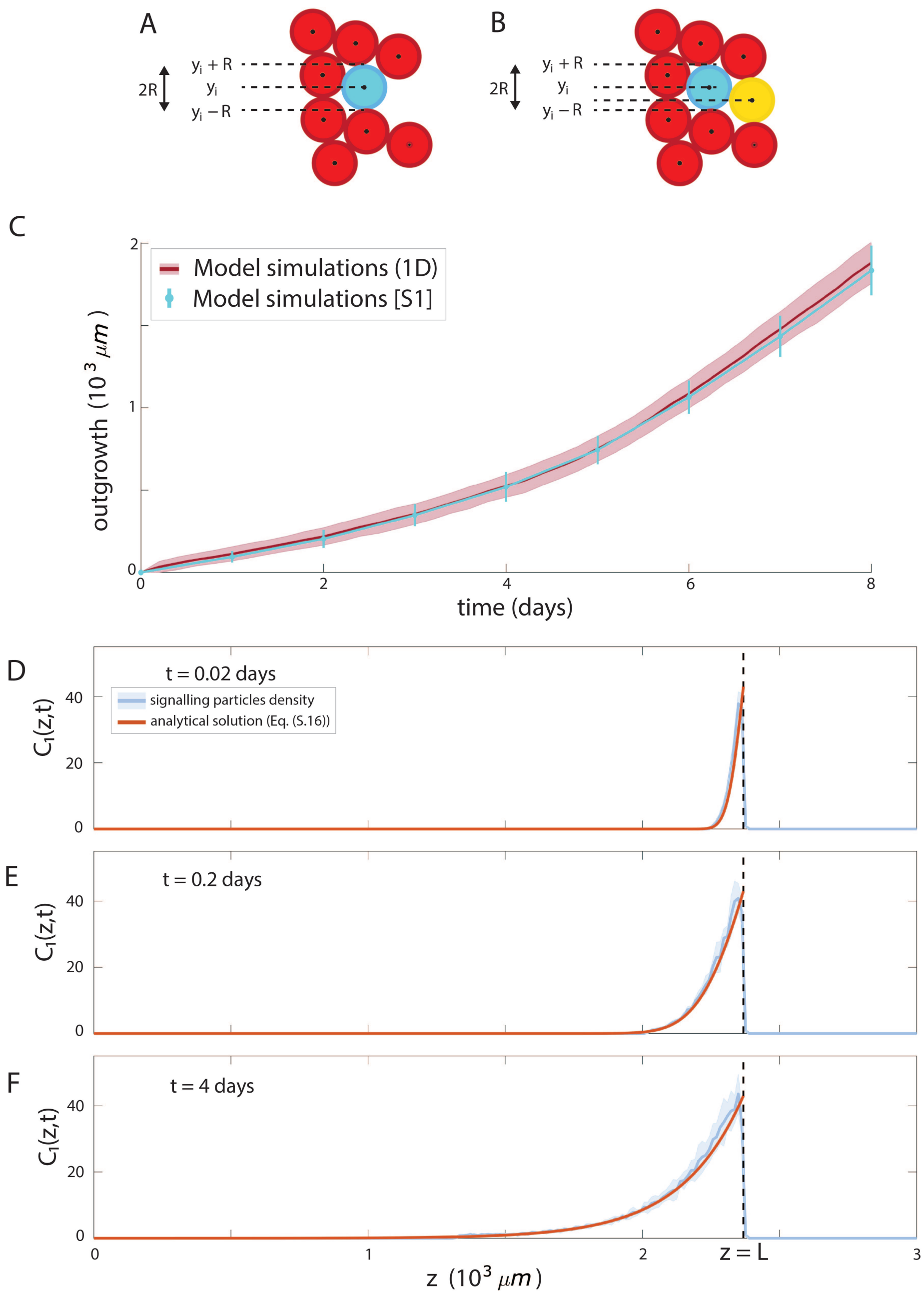


Figure S1: **Validation of the spatial scale of the model, Related to Figure 2 and Section 2.1 of the main text (A,B)** Scheme showcasing two cases where ependymal cell $i$ (cyan disc) is a front cell (panel A) or not (panel B), using the identification rule described in section S2.1. **(C)** Comparison between the outgrowth predicted by our simplified 1D model (red shadowed area) and by[1] (cyan error bar). Each prediction has been obtained averaging over 100 replicates. Means are represented as lines, shadowed area and error bar correspond to 1 standard deviation. **(D,E,F)** Comparison between the 1D density profile of signalling particles (both the mean - blue solid line - and the standard deviation - blue shaded area - are calculated from the $y$-axis at each $x$ position) with the analytical solution $C_1(z,t)$ (Eq. (S.11)) of the reaction-diffusion model in finite domain (orange solid line). For $D = 0.6\ \mu m^2 \cdot s^{-1}$, $k = 1$ days$^{-1}$, at different times (from top to bottom: day 0.02, day 0.2, day 4).

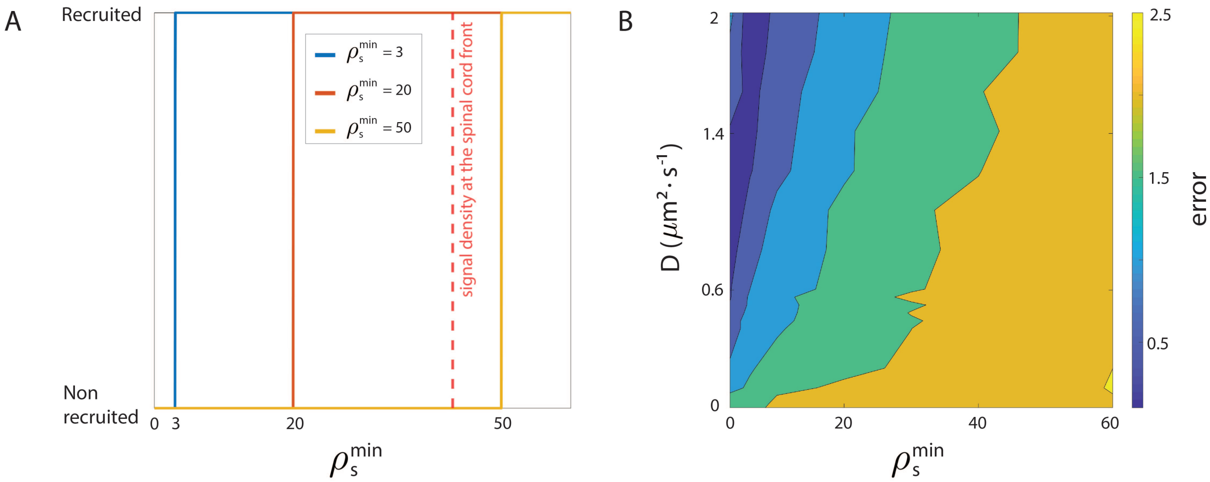


Figure S2: **Role of the cell sensitivity to the signal density, Related to Figure 3 and Section 2.3 of the main text. (A)** Cell activation function as function of the local sensitivity of the ependymal cells, for different sensitivity levels to the signal (parameter $\rho_s^{min}$). The maximal signal density (imposed on the right boundary, i.e. on the spinal cord front) is indicated in red. For small sensitivity to the signal (large $\rho_s^{min}$, yellow curve), no cells will be recruited by the signal, while for large sensitivity (small $\rho_s^{min}$, blue curve), the majority of cells exposed to the signal will be recruited instantaneously. **(B)** Error between the model predicted recruitment limit and experimental recruitment limit curve [5] as function of the sensitivity to the signal $\rho_s^{min}$ (x-axis) and the diffusion coefficient $D$ (y-axis).

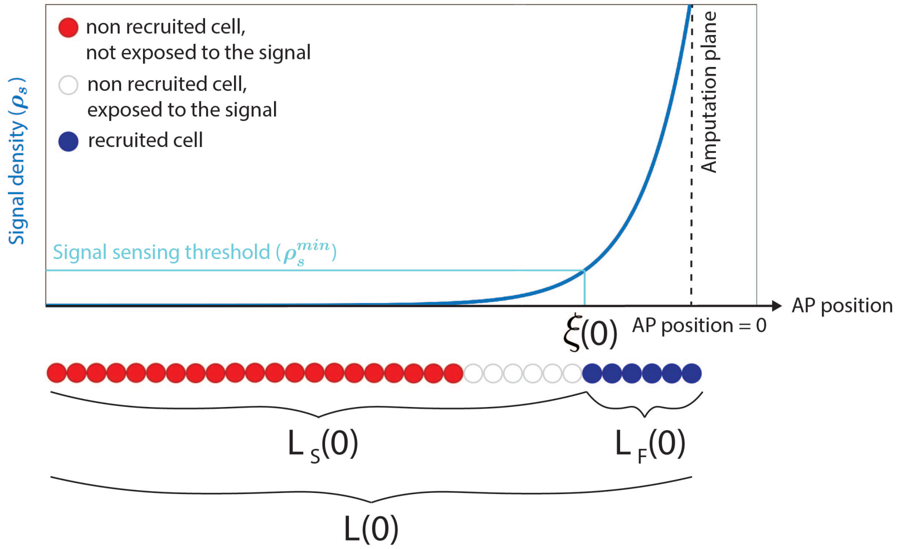


Figure S3: **Theoretical prediction of tissue outgrowth, Related to Figure 4 and Section 2.4 of the main text**. Tissue domain occupied by recruited fast proliferating cells ($L_F(0)$), tissue domain occupied by non-recruited slow proliferating cells ($L_S(0)$) and recruitment limit ($\xi(0)$) at time 0. The x-axis represents the AP axis, the blue curve is the density of the signal. The amputation plane is indicated by a dashed black line. The signal sensing threshold ($\rho_s^{min}$) for cell recruitment is indicated in cyan.

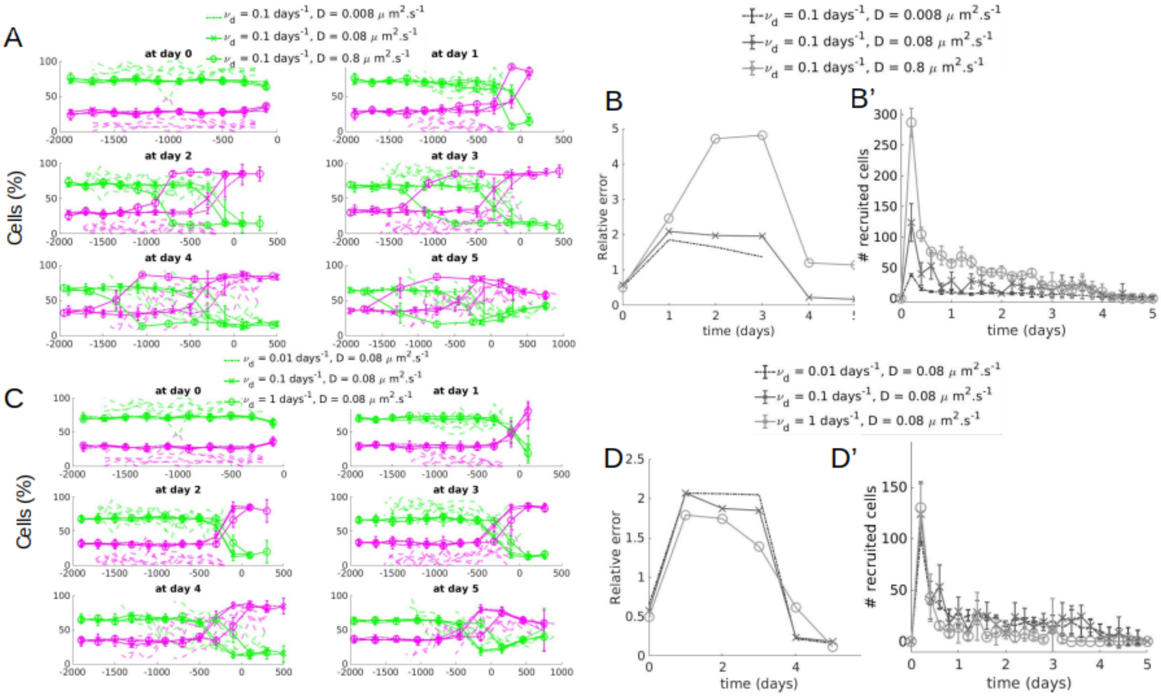


Figure S4: **Signal-dependent transient modulation of cell cycling, Related to Figure 5 and Section 2.5 of the main text. (A, C)** Percentage of cells along the AP-axis that are in G0/G1 phase (green curves) and in S/G2 phase (magenta) as function of the AP position for different times (different subplots). Dotted lines are experimental data from[1], solid lines are simulations for different values of $D$ and fixed $\nu_d = 0.1 days^{-1}$ (A) or different values of $\nu_d$ and fixed $D = 0.08\mu m^2.s^{-1}$ (C). For each set of parameters, simulated data are averaged over 10 realisations (solid lines represent the mean, errorbars the standard deviation). **(B, D)** Relative error (see section 8 for details on its computation) between the simulated and experimental cell distributions as function of time post-amputation for different values of $D$ while fixing $\nu_d = 0.1 days^{-1}$ (B), and for different values of $\nu_d$ while fixing $D = 0.08\mu m^2.s^{-1}$(D). **(B', D')** Number of newly recruited cells by the signal computed every 0.2 days as a function of time for different values of $D$ while fixing $\nu_d = 0.1 days^{-1}$ (B'), and for different values of $\nu_d$ while fixing $D = 0.08\mu m^2.s^{-1}$(D').

| Name | Value (range) | Description |
|---|---|---|
| $D$ | $[0, 2]\ \mu m^2 \cdot s^{-1}$ | Signal diffusion constant |
| $k$ | $[0.5, 100]\ day^{-1}$ | Degradation rate |
| $\rho_0$ | 42.98 cell.$(13\mu m)^{-2}$ | Signal density imposed at the front (Dirichlet condition) |
| $\rho_s^{min}$ | $[3, 60]$ cell.$(13\mu m)^{-2}$ | Density threshold for cell recruitment |
| $N_0$ | 1676 | Initial number of cells |
| $R$ | 6.6 $\mu$m | Cell radius |
| $\mu_S$, $\sigma_S$ | 340 h $\pm$ 32h | Cell cycle length for slow cycling cells[2] |
| $\mu_F, \sigma_F$ | 119h $\pm$ 10h | Cell cycle length for fast cycling cells[2] |
| $L_0$ | 2.36 mm | Initial tissue length (x-direction, AP axis) |
| $\{y_{\min}, y_{\max}\}$ | $\{-52.8, 52.8\}\mu m$ | lower and upper limits in y-direction |

Table S1: **Model parameters, Related to STAR method section of the main text (section Mathematical modelling)**. Model parameters and experimental values.

### 2 Supplementary controls

#### 2.1 Validation of the modelling of the spatial motion of the ependymal cells

In this section, we aim to validate the cellular scale of our model (described in the STAR Methods section of the main text) by using the 1D model developed in[1] for spinal cord regeneration. Indeed, the model of[1] bears analogies with our own model as cell division follows the same rules as described in the

STAR Methods section of the main text and cells are supposed to behave as rigid spheres. However, cell motion is treated differently in[1] as upon division, one daughter cell occupies the same position of the mother cell and the other daughter cell is placed posteriorly to the mother cell, moving all the posterior cells a quantity corresponding to the size of a cell in the posterior direction. This corresponds exactly to the dynamics of rigid spheres in a 1D setting, but does not extend easily to 2D, justifying the need for the more complex treatment of cell motion that we proposed (see STAR Methods section of the main text). The goal of this section is therefore to validate our treatment of cell motion by comparing it to the exact motion of rigid spheres provided in[1].

To this aim, we also need to adapt our signalling process, as in[1] the ependymal cells are supposed to be recruited by a signal spreading with constant velocity from the amputation plane, contrary to our more realistic model of a signal evolving via a reaction diffusion equation (see STAR Methods section of the main text). Therefore in this section, we consider the same signal as in[1], i.e a signal moving with constant velocity from the amputation plane, during $\tau = 85 \pm 12$ hours and recruiting ependymal cells within $\lambda = 828 \pm 30$ $\mu$m from the amputation plane.

In Fig. S1 C, we show the tissue outgrowth (distance from the amputation plane to the the most posterior ependymal cell) obtained with our modified model in 1D (red curve) compared to the outgrowth of with the same set of parameters as in[1]. Our model prediction is in good agreement with the previous results[1], validating the cellular scale of our model in 1D.

#### 2.2 Validation of the numerical scheme for the reaction-diffusion process of the signal

In this section, we aim to validate the numerical scheme of the reaction-diffusion process governing the signaling scale of our model (see STAR Methods section of the main text). To this aim, we consider a simplified model where the tissue is fixed (no cell division), and 1D. Under these conditions, the equation for the signal simplifies to a reaction-diffusion equation on a fixed 1D domain with Dirichlet boundary conditions on one side and no-flux boundary conditions on the other side. This enables to derive an analytical solution adapting the method used in**ceccarelli2021** to our setting, and compare our numerical simulations with the exact solution obtained.

*Derivation of the analytical solution of the reaction-diffusion model in finite domain.*

Here, we derive an analytical solution for a reaction-diffusion equation in a finite and fixed 1D domain, with non-homogeneous Dirichlet conditions on one side and Neumann boundary conditions (null flux) on the other side. We adapt the derivation of**ceccarelli2021**, where the authors considered various but different boundary conditions.

We consider a signal spreading in a non-growing 1D tissue of length $L$ and we call $C_1(z,t)$ its concentration, with $z \in [0, L]$ and $t \geq 0$. In order to adapt the derivation of**ceccarelli2021**, we consider a new spatial variable $x = L - z$ and a 1D tissue of length $L$ starting at $x = 0$. We assume that the signal concentration is given at $x = 0$, it diffuses to the tip of the tissue with a diffusion constant $D$ and degrades linearly at a rate $k$. We assume no-flux boundary conditions at the tip of the tissue in $x = L$. At $t = 0$ there is no signal in the tissue. The evolution in space and time of the signal distribution $C_1$ is described by the following reaction-diffusion equation

$$\frac{\partial C_1}{\partial t} = D\frac{\partial^2 C_1}{\partial x^2} - kC_1. \tag{S.1}$$

We have the following initial and boundary conditions:

- no signal at initial time:
$$C_1(x, t = 0) = 0,$$
- fixed signal concentration at $x = 0$:
$$C_1(x = 0, t) = C_0,$$
where $C_0$ is constant,

- Neumann boundary conditions at the tip of the tissue $x = L$:

$$\frac{\partial C_1}{\partial x}(x = L, t) = 0.$$

We introduce the dimensionless variables

$$\varepsilon = \frac{x}{\lambda} \qquad \text{and} \qquad \tau = kt, \tag{S.2}$$

where $\lambda = \sqrt{\frac{D}{k}}$ is the characteristic reaction-diffusion length, and we rewrite Eq. (S.1). We define the quantity $\ell = \frac{L}{\lambda}$, which is now the only model parameter. We rewrite the concentration as $C(\varepsilon, \tau) = \frac{C_1(\varepsilon,\tau)}{C_0}$ and we obtain the following equation

$$\frac{\partial C}{\partial \tau} = \frac{\partial^2 C}{\partial \varepsilon^2} - C \tag{S.3}$$

with the following initial and boundary conditions:

- no morphogen at initial time:

$$C(\varepsilon, \tau = 0) = 0,$$

- fixed signal concentration at $\varepsilon = 0$:

$$C(\varepsilon = 0, \tau) = 1,$$

  where $C_0$ is constant,

- Neumann boundary conditions at the tip of the tissue $\varepsilon = \ell$:

$$\frac{\partial C}{\partial \varepsilon}(\varepsilon = \ell, \tau) = 0.$$

To solve this equation we introduce an auxiliary function $C_2$ and we define $C$ as follows:

$$C = C_2 e^{-\tau}.$$

In order to rewrite Eq. (S.3) in terms of $C_2$, we compute the spatial and time derivatives of $C$ as follows:

$$\frac{\partial C}{\partial \varepsilon} = \frac{\partial C_2}{\partial \varepsilon} e^{-\tau} \qquad \Rightarrow \qquad \frac{\partial^2 C}{\partial \varepsilon^2} = \frac{\partial^2 C_2}{\partial \varepsilon^2} e^{-\tau}, \tag{S.4}$$

$$\frac{\partial C}{\partial \tau} = e^{-\tau} \frac{\partial C_2}{\partial \tau} - C_2 e^{-\tau}. \tag{S.5}$$

Inserting $\frac{\partial^2 C}{\partial \varepsilon^2}$ and $\frac{\partial C}{\partial \tau}$ computed in Eq. (S.4) and Eq. (S.5), respectively, into Eq. (S.3), leads to the following equation

$$\frac{\partial C_2}{\partial \tau} = \frac{\partial^2 C_2}{\partial \varepsilon^2} \tag{S.6}$$

with the following conditions

$$C_2(\varepsilon, \tau = 0) = 0,$$
$$C_2(\varepsilon = 0, \tau) = e^{\tau},$$
$$\frac{\partial C_2}{\partial \varepsilon}(\varepsilon = \ell, \tau) = 0.$$

To solve Eq. (S.6) we have to redefine $C_2$ using the auxiliary functions $C_2(\varepsilon, \tau)$, $f(\tau)$ and $g(\varepsilon)$. We define $C_2(\varepsilon, \tau)$ as

$$C_2(\varepsilon, \tau) = C_3(\varepsilon, \tau) + f(\tau) g(\varepsilon),$$

while $f(\tau)$ and $g(\varepsilon)$ will be defined explicitly later on. The time derivative of $C_2(\varepsilon, \tau)$ is

$$\frac{\partial C_2}{\partial \tau}(\varepsilon, \tau) = \frac{\partial C_3}{\partial \tau}(\varepsilon, \tau) + g(\varepsilon)\frac{\partial f}{\partial \tau}(\tau) \tag{S.7}$$

and the spatial derivative is

$$\frac{\partial^2 C_2}{\partial \varepsilon^2}(\varepsilon, \tau) = \frac{\partial^2 C_3}{\partial \varepsilon^2}(\varepsilon, \tau) + \frac{\partial^2 g}{\partial \varepsilon^2}(\varepsilon)f(\tau). \tag{S.8}$$

Again, inserting $\frac{\partial C_2}{\partial \tau}$ and $\frac{\partial^2 C_2}{\partial \varepsilon^2}$ computed in Eq. (S.7) and Eq. (S.8), respectively, into Eq. (S.6), allows to rewrite the reaction-diffusion equation in $C_3$ as

$$\frac{\partial C_3}{\partial \tau}(\varepsilon, \tau) = \frac{\partial^2 C_3}{\partial \varepsilon^2}(\varepsilon, \tau) + \frac{\partial^2 g}{\partial \varepsilon^2}(\varepsilon)f(\tau) - g(\varepsilon)\frac{\partial f}{\partial \tau}(\tau) \tag{S.9}$$

with the following conditions:

- the initial condition:

$$C_3(\varepsilon, \tau = 0) = -g(\varepsilon)f(\tau = 0),$$

- the source:

$$C_3(\varepsilon = 0, \tau) = e^{\tau} - g(\varepsilon = 0)f(\tau),$$

- the Neumann boundary conditions:

$$\frac{\partial C_3}{\partial \varepsilon}(\varepsilon = \ell, \tau) = -\frac{\partial g}{\partial \varepsilon}(\varepsilon = \ell)f(\tau).$$

It is preferable to have the initial condition different from 0 and the boundary conditions equal to 0. Hence, we define $g(\varepsilon)$ and $f(\tau)$ as:

$$g(\varepsilon) = 1 \qquad \text{and} \qquad f(\tau) = e^{\tau}.$$

Then, Eq. (S.9) becomes

$$\frac{\partial C_3}{\partial \tau}(\varepsilon, \tau) = \frac{\partial^2 C_3}{\partial \varepsilon^2}(\varepsilon, \tau) - e^{\tau} \tag{S.10}$$

with the following conditions

- the initial condition:

$$C_3(\varepsilon, \tau = 0) = -1 \neq 0,$$

- the source:

$$C_3(\varepsilon = 0, \tau) = 0,$$

- the Neumann boundary conditions:

$$\frac{\partial C_3}{\partial \varepsilon}(\varepsilon = \ell, \tau) = 0.$$

The solution to systems of the type of Eq. (S.10) can be found in[3]. The solution to the problem will be

$$u(x,t) = \sum_{j=0}^{\infty} f_j v_j(x) e^{-\lambda_j \int_0^t \frac{1}{m(s)} ds} + \sum_{j=0}^{\infty} v_j(x) e^{-\lambda_j \int_0^t \frac{1}{m(s)} ds} \int_0^t \frac{F_j(s)}{m(s)} e^{-\lambda_j \int_0^s \frac{1}{m(w)} dw}.$$

Following what the authors have done in**ceccarelli2021** we find that

$$v_j(\varepsilon) = \sqrt{\frac{2}{\ell}} \sin\left(\frac{(j+\frac{1}{2})\pi}{\ell}\varepsilon\right),$$

$$\lambda_j = \left(\frac{(j+\frac{1}{2})\pi}{\ell}\right)^2,$$

$$f_j = -\frac{1}{\sqrt{\lambda_j}}\sqrt{\frac{2}{\ell}},$$

$$F_j(\tau) = -\frac{e^{\tau}}{\sqrt{\lambda_j}}\sqrt{\frac{2}{\ell}}.$$

Hence

$$C_3(\varepsilon,\tau) = \sum_{j=0}^{\infty} -\frac{1}{\sqrt{\lambda_j}}\frac{2}{\ell}\sin(\sqrt{\lambda_j}\varepsilon)e^{-\lambda_j\tau}\left(\frac{\lambda_j + e^{-(1+\lambda_j)\tau}}{1+\lambda_j}\right).$$

Then, since

$$C_2(\varepsilon,\tau) = C_3(\varepsilon,\tau) = f(\tau)g(\varepsilon),$$
$$C(\varepsilon,\tau) = C_2(\varepsilon,\tau)e^{-\tau} = C_3(\varepsilon,\tau)e^{-\tau} + 1,$$

finally we have

$$C(\varepsilon,\tau) = \sum_{j=0}^{\infty} -\frac{1}{\sqrt{\lambda_j}}\frac{2}{\ell}\sin(\sqrt{\lambda_j}\varepsilon)e^{-\lambda_j\tau}\left(\frac{\lambda_j e^{-(1+\lambda_j)\tau} + 1}{1+\lambda_j}\right) + 1.$$

Finally, using (S.2) and the change of variables $x = L - z$, we write the concentration $C_1$ as

$$C_1(z,t) = C_0\, C\left(\frac{L-z}{\lambda}, t\right). \tag{S.11}$$

*Numerical validation*

In order to solve numerically the reaction-diffusion equation ( Eq. (6) of the STAR Methods section of the main text), we discretize the continuous signal density $\rho_S(x,t)$ using the Smoothed Particle Hydrodynamic (SPH) method. As explained in the STAR Methods section of the main text, we passed to an agent-based model by approximating the continuous density $\rho_S(x,t)$ with a set of discrete agents called signaling particles (SPs) that are continuously produced at the tip of the tissue, diffuse through the tissue with speed $D$ and are removed randomly from the domain according to a Poisson process of rate $k$.

Considering a finite domain consisting of non-proliferating ependymal cells, we compare the 1D density profile of SPs with the analytical solution given by Eq. (S.11). In Figure S1 D, E, F, we observe a good agreement at different times between the numerical and analytical solution of the reaction-diffusion model in finite domain. This allows to validate the numerical scheme for the reaction-diffusion process of the SPs.
